# Proposing a methodology for axon-centric analysis of IOP-induced mechanical insult

**DOI:** 10.1101/2024.09.09.611631

**Authors:** Manik Bansal, Bingrui Wang, Susannah Waxman, Fuqiang Zhong, Yi Hua, Yuankai Lu, Juan Reynaud, Brad Fortune, Ian A. Sigal

## Abstract

**Purpose:** IOP-induced mechanical insult on retinal ganglion cell axons within the optic nerve head (ONH) is believed to be a key factor in axonal damage and glaucoma. However, most studies focus on tissue-level mechanical deformations, overlooking that axons are long and thin, and that their susceptibility to damage likely depends on the insult’s type (e.g. stretch/compression) and orientation (longitudinal/transverse). We propose an axon-centric approach to quantify IOP-induced mechanical insult from an axon perspective.

**Methods:** We used optical coherence tomography (OCT) scans from a healthy monkey eye along with histological images of cryosections to reconstruct the axon-occupied volume including detailed lamina cribrosa (LC) pores. Tissue-level strains were determined experimentally using digital volume correlation from OCT scans at baseline and elevated IOPs, then transformed into axonal strains using axon paths estimated by a fluid mechanics simulation.

**Results:** Axons in the LC and post-LC regions predominantly experienced longitudinal compression and transverse stretch, whereas those in the pre-LC and ONH rim mainly suffered longitudinal stretch and transverse compression. No clear patterns were observed for tissue-level strains.

**Conclusions:** Our approach allowed discerning axonal longitudinal and transverse mechanical insults, which are likely associated with different mechanisms of axonal damage. The technique also enabled quantifying insult along individual axon paths, providing a novel link relating the retinal nerve fiber layer and the optic nerve through the LC via individual axons. This is a promising approach to establish a clearer connection between IOP-induced insult and glaucoma. Further studies should evaluate a larger cohort.

## 1. Introduction

The optic nerve head (ONH) is the initial site of retinal ganglion cell (RGC) damage in glaucoma, particularly within the lamina cribrosa (LC) region ^1, 2^. While the exact causes of RGC damage in glaucoma are not fully understood, there is a strong association between the onset and progression of glaucoma and elevated intraocular pressure (IOP) ^3, 4^. Furthermore, all currently approved treatments for glaucoma are based on lowering IOP ^5^. The role of IOP on glaucoma, however, is complicated ^4, 6, 7^. Some patients suffer glaucoma at statistically normal levels of IOP, whereas others with elevated IOP do not have clear pathology or progress much slower ^3, 4^. This complex relationship between IOP and glaucomatous neuropathy suggests that susceptibility to glaucoma is more closely determined by IOP-induced tissue deformations, than by the direct level of IOP alone ^8^.

Vision loss in glaucoma is irreversible due to the current inability to regenerate RGCs ^9, 10^. Understanding of the mechanisms through which IOP causes or contributes to damage RGC axons is crucial for improving preventive strategies against vision loss. Experimental ^11–15^ and computational ^16–18^ techniques have been used to measure and predict the tissue deformations caused by the elevated IOP, aiming to understand their relationship with the glaucomatous neural tissue loss. These deformations are multidimensional, involving stretch, compression, and shear simultaneously (in different directions)^6, 12^, which has made the analysis and interpretation of the many outcomes of the studies quite complex. In an attempt to simplify the analysis, some authors have suggested using equivalent strain as a single measure of deformation ^19^. Whilst the results are indeed simpler when looking at a single outcome measure, the studies using the simplifications have not fared any better than the studies using many measures in identifying a clear relationship between insult and glaucoma. We reason that different IOP-induced deformations are likely to contribute to RGC axon damage through different mechanisms. Hence, we propose that a step forward will result from evaluating the effects of IOP from an axonal perspective that distinguishes the different types of insult.

**We postulate that to understand how IOP affects axons and contributes to axonopathy, it is essential to investigate the effects of IOP from the axon perspective.** This requires investigating beyond the general “bulk” tissue-level deformations that other studies have measured or predicted, and focusing on how these distortions affect the axon directly. The geometric features of RGC axons such as their length (50-100 mm in primates), the nearly 90° turn upon transitioning from the retinal nerve fiber layer (RNFL) to the ON, and the passing through narrow LC pores are thought to contribute to heightened vulnerability to damage ^20, 21^. These features of axon paths suggest that the axons vary in their vulnerability to damage depending on what the insult looks like from the perspective of the axon. For instance, compression transverse to an axon may directly disrupt axonal transport and axoplasmic flow, whereas longitudinal compression may cause axonal bending or buckling ^10, 22–24^. The two insults thus may affect axons differently, and at different magnitudes, with different time scales and potentially different abilities to recover.

Our goal was to describe and demonstrate an axon-centric approach for quantifying the in vivo mechanical insult to RGC axons within the ONH under elevated IOP. Based on the optical coherence tomography (OCT) scans of a healthy monkey eye obtained at controlled IOPs (baseline and elevated), we first determined the IOP-induced “bulk” tissue deformations. We approximated the 3D axon paths within the ONH, from the RNFL, through the LC and to the optic nerve. The experimentally-determined IOP-induced tissue-level deformations were then transformed into axonal deformations, separating longitudinal and transverse distortions to axons. To evaluate potential advantages of the axon-centric approach, we analyzed the distributions of the axonal insults. We thus demonstrate how the integration of experimental and computational techniques can facilitate a deeper understanding of axon damage mechanisms.

## 2. Methods

Our goal was to demonstrate an axon-centric approach for quantifying the mechanical insult from the axon perspective within the ONH under elevated IOP. The eyes of an adult female rhesus macaque monkey (*Macaca mulatta*) were used as a proof of concept in this study. Our general strategy involved the following main steps (**Figure 1**):

1. in vivo and histological imaging of the ONH: Optical coherence tomography (OCT) was used for in vivo imaging of ONH at different IOP levels, whereas the cryosectioned ONH was imaged with instant polarized light microscopy (IPOL).
2. Quantification of tissue-level mechanical insult resulting from an IOP increase: Using digital volume correlation (DVC) to measure deformations from OCT images acquired at baseline (IOP = 10 mm Hg) and elevated (IOP = 40 mm Hg) IOPs.
3. Reconstruction of a 3D eye-specific ONH volume for the axons: The reconstruction is done from in vivo OCT scans, supplemented by data from histology for the LC beam structures that are not well visualized in OCT.
4. Estimation of the axon paths: Using a fluid mechanical simulation within the non-collagenous axonal volume, the axon paths were estimated for the specific monkey eye.
5. Quantification of the axonal mechanical insult: Integrate the tissue-level mechanical insult with the estimated axon paths to quantify the mechanical insult longitudinally and transverse to the axons.

**Figure 1:**
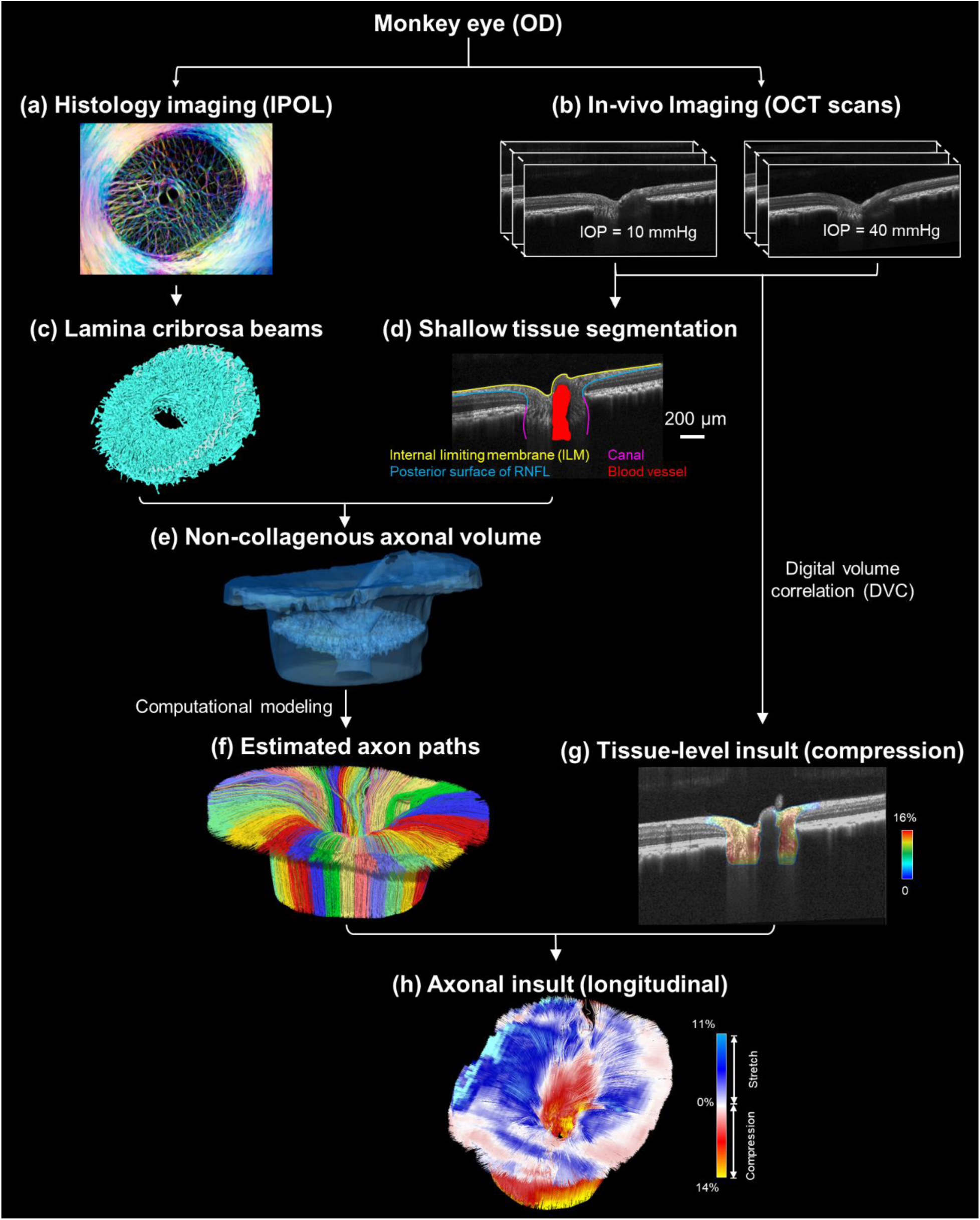
Workflow of the proposed axon-centric approach: (1) in vivo and histological imaging of the ONH: Optical coherence tomography (OCT) was used for in vivo imaging of ONH at different IOP levels, while the sectioned ONH was imaged with Instant polarized light microscopy (IPOL). (2) Quantification of tissue-level mechanical insult resulting from an IOP increase: Using digital volume correlation (DVC) to measure deformations from OCT images acquired at baseline (IOP = 10 mm Hg) and elevated (IOP = 40 mm Hg) IOPs. (3) Reconstruction of a 3D eye-specific ONH volume for the axons: The 3D shallow tissue was segmented based on the OCT scans at the baseline IOP, while the LC beam structure was reconstructed from the IPOL images. Removing the LC beams structure from the shallow tissue volume can obtain the non-collagenous axonal volume. (4) Estimation of the axon paths: Using a fluid mechanical simulation within the non-collagenous axonal volume, the axon paths were estimated for the specific monkey eye. (5) Quantification of the axonal mechanical insult: Integrating the tissue-level mechanical insult with the estimated axon paths, the mechanical insults longitudinal and transverse to each axon were obtained.

Each of the steps is described in detail below.

### 2.1 In-vivo and histological imaging of ONH

All procedures were carried out in strict accordance with the recommendations in the Guide for the Care and Use of Laboratory Animals of the National Institutes of Health and were approved and monitored by the Institutional Animal Care and Use Committee (IACUC) at Legacy Health (US Department of Agriculture license 92-R-0002; OLAW assurance A3234-01; AAALAC International assurance #000992). All live animal procedures also adhered to the Association for Research in Vision and Ophthalmology’s Statement for the Use of Animals in Ophthalmic and Vision Research.

#### In vivo imaging of ONH (OCT scans)

The in vivo experiment procedure followed an approach similar to the one described elsewhere ^25^. Briefly, induction of general anesthesia was followed by intubation with an endotracheal tube to breathe a mixture of medical air and oxygen. To sustain anesthesia throughout the session, 0.75-2.0% isoflurane was added to the inspired gas. Vital signs were monitored throughout each imaging session and recorded every 15 min. A commercial OCT device (SPECTRALIS OCT2 + HRA, SPX1902 version 6.12, Heidelberg Engineering, GmbH, Heidelberg, Germany) was used to scan the ONH in-vivo under different IOP conditions. A super luminescence diode source with 870 nm central wavelength and 50 nm bandwidth was used in the device during scanning. A primary difference with the previous study was the OCT scan pattern. For the OCT imaging in this work we acquired a dense isotropic grid scan pattern comprised of 768 x 768 A-lines. This scan covered a total area of 15° x 15° centered on the ONH, with the real time averaging set to 4x. Follow-up OCT scans were acquired at the same location as the baseline scan using the instrument’s automatic active eye tracking software. After acquisition, all OCT scans were exported in *.vol format for DVC calculation and reconstruction of a 3D eye-specific ONH volume ^25^. Note that the eye used in this study had also been used to study longitudinal experimental glaucoma. However, we only used the OCT data from when the eye was healthy.

#### Histology imaging of the LC

Some morphological details of the LC beam structure were not discernible in OCT scans due to limitations of OCT imaging, particularly for the deeper portions of the canal where OCT signal is weakest. We therefore supplemented the canal architecture using 3D LC beam information from histology. The process was as follows: Both eyes of the monkey scanned with OCT were perfusion fixed at an IOP of 10 mmHg at the time of sacrifice and then enucleated. The reconstruction after enucleation followed the general approach for collagen beams described elsewhere, except that no vessel labels were used ^26^. Briefly, the ONH was isolated using a 14-mm-diameter trephine. The ONH was cryoprotected in a 30% sucrose solution overnight, flash-frozen in optimum cutting temperature compound (Tissue Plus, Fisher Health-care, Houston, TX), and sectioned coronally at 16 μm thickness with a cryostat (Leica CM3050S). IPOL was implemented using a 4× strain-free objective (UPLFLN 4XP, Olympus, Tokyo, Japan) to visualize LC collagen beams ^27^. Sequential IPOL images were aligned to generate image stacks, which were later used for the reconstruction of LC beams and surrounding sclera and pia mater ^26, 27^. The IPOL images have high spatial resolution and reliably measure the LC architecture without the problem of shadows from blood or signal decay with depth normally present in OCT scans of the living eye.

### 2.2 Quantification of tissue-level mechanical insult resulting from an IOP increase

We utilized the digital volume correlation (DVC) technique to evaluate the bulk or tissue-level acute IOP-induced deformations in the ONH, including tissue-level stretch, compression and shear (**Fig. 1**). This DVC methodology was specially developed for the same set of OCT images. To improve the accuracy, we used 1) a pre-registration technique to remove large ONH rigid body motion in OCT volumes, 2) a modified 3D inverse-compositional Gaussian Newton method to ensure sub-voxel accuracy of displacement calculations despite high noise and low image contrast of some OCT volumes, and 3) a confidence-weighted strain calculation method was applied to further improve the accuracy. We have tested that our DVC method had displacement errors smaller than 0.037 and 0.028 voxels with Gaussian and speckle noises, respectively. The strain errors in the three directions were less than 0.45% and 0.18% with Gaussian and speckle noises, respectively. The details of the tests that led to these conclusions are all described in Zhong 2022 ^25^.

The analysis was based on images acquired at baseline 10mmHg and elevated 40mmHg IOPs. The OCT volume at 10 mm Hg was considered as reference (undeformed) set of images, while that at 40 mmHg was considered the deformed set of images. The detailed DVC process was described previously in detail ^25^. Briefly, this procedure involves three steps: 1) pre-processing of OCT scans (scaling, making isotropic, gross solid-body registration, etc.); 2) displacement measurement by correlation of reference and deformed OCT volumes; and 3) strain calculation at each pixel by evaluating the deformation gradients. The resultant strain field was strain tensors in a Cartesian coordinate system, with the x-axis along the inferior-superior, the y-axis along the nasal-temporal, and the z-axis along the anterior-posterior direction.

### 2.3 Reconstruction of a 3D eye-specific ONH volume for the axons

The 3D eye-specific non-collagenous axonal volume was reconstructed in three steps: 1) Shallow tissues of ONH volume were reconstructed from OCT images, 2) Deep tissues i.e., detailed beam microstructure of LC (to identify the LC pores), were reconstructed from IPOL images, and 3) Integration of shallow and deep tissue volumes to obtain a non-collagenous axonal volume. Both OCT and histology imaging for the reconstruction were obtained at baseline IOP of 10 mmHg. Each step is detailed below.

#### Shallow tissue volume reconstruction

The shallow tissue volume was reconstructed by segmenting visible portions in the OCT scans obtained at IOP of 10 mm Hg. Image segmentation was performed manually using Avizo (version 9.1, FEI; Thermo Fisher Scientific). The shallow tissue volume was first delineated within the bounds of three image features: the internal limiting membrane (ILM), the posterior surface of RNFL, and the canal. To minimize morphometric errors, only the visible region near the periphery of the canal were segmented. Then the central retinal vessels (CRVs) and their bifurcations were marked and excluded from the segmented volume. In all OCT scans, the shadow in the canal region was marked as the central CRVs. The marked CRV region was then checked and corrected to match the location of CRVs in the C-scan images. To avoid divergent effects at the corners of a rectangular boundary, we set an elliptical boundary for the RNFL.

#### Deep tissue volume reconstruction

The model reconstruction demonstrated in this work was based on OCT scans obtained from a healthy normal eye at an IOP of 10mmHg. The LC beam structure from histology used to supplement the OCT data was from the contralateral control eye, mirrored in the nasal/temporal direction and carefully aligned. The rationale for using the contralateral eye and potential consequences is addressed in the Discussion.

#### Integration of shallow and deep tissue volume

To integrate the shallow and deep tissue volumes, the deep tissue volume was positioned at the LC location within the shallow tissue volume. It was mirrored in the nasal-temporal direction, and translated and rotated for alignment between the CRVs and the optic canal periphery. The process of alignment of LC between the shallow and deep volumes involves the following steps: 1) Delineating the anterior LC surface in the OCT volume. We have already demonstrated elsewhere that we can do this with very high reliability and reproducibility. ^28–30^ 2) We identified landmarks at the superior, inferior, nasal, and temporal points on both the anterior LC surfaces in both the shallow and deep volumes. 3) We calculated the transformation that brings the two sets of landmarks into coincidence. 4) The calculated transformation was applied to the deep tissue volume to align with the shallow tissue volume. The alignment of the deep and shallow tissues is illustrated in **Figure 2**. The aligned deep tissue volume was then removed from the shallow tissue volume to obtain the non-collagenous axonal volume.

**Figure 2.**
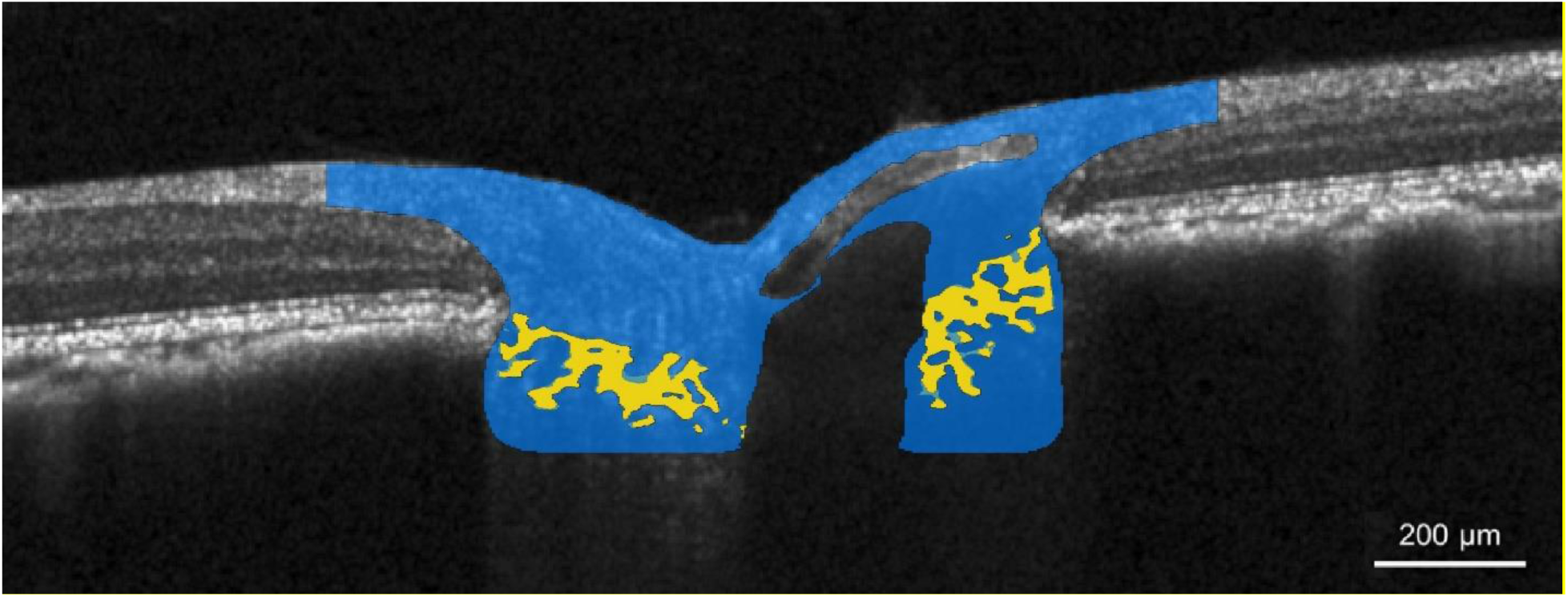
Alignment of deep and shallow tissue volumes. The blue shading indicates the region of the tissue volume in which axons are allowed, from the RNFL layer at the periphery to the ON. The yellow shadowing indicates the LC beams obtained from the deep tissue volume from histology, overlaid on the full tissue volume.

### 2.4 Estimation of the axon paths

We assumed that the axon paths from the RNFL to the optic nerve are smooth and continuous. They will not pass through the LC beams, the CRVs or themselves (i.e. no crossings, splits, bifurcations or similar). We recognized that these assumptions are similar to those made for fluid mechanics. We thus solved a fluid mechanics simulation within the non-collagenous axonal volume. The volumetric mesh of the non-collagenous axonal volume was generated in Avizo with 19.4 million tetrahedral elements (**Supplementary Figure 3**). The fluid within the non-collagenous axonal volume was modeled as an incompressible Newtonian fluid with a density of 1000 kg/m^3^ and a viscosity of 0.002 Pa s. Boundary conditions for this simulation were applied: 1) inlet on the bottom surface of ONH, with a pressure of 10 kPa. 2) outlet on the elliptical boundary surface of RNFL, with zero pressure. 3) non-slip boundary conditions for other surfaces. The simulation only considered laminar flows, with Reynolds number of the flow in the range of 1–100.

After the flow was simulated, to trace the axon paths within the ONH, streamlines were generated from the velocity field resulting from the fluid mechanical simulation. Streamlines are defined as the paths of massless particles moving with the flow, providing an ideal reference for tracing axon paths within the region from RNFL to the optic nerve (**Figure 3**). The tracing was started at the inlet surface, with each streamline following the direction of the velocity vector at sequential points to replicate a potential axon path. Because of approximations made during integration, some streamlines appeared to end at the boundary, which was inconsistent with our assumption of axonal continuity. This could be solved by reducing the integration step, but this, in turn, substantially increased the time needed to follow the streamlines, and the computational requirements necessary to store, process and visualize them. Instead, we excluded streamlines that did not reach between the boundaries or were discontinuous. In this specific monkey eye, a total of 31815 axon paths were traced, with each axon delineated by a varying number of points, ranging from 240 to 617 (one example axon path is demonstrated in **Figure 4a**). The number of points depended on the complexity of the path. The simplest path, a straight line, can be represented by just two points. More complex paths required more points. It is equally possible to trace a larger number of axons, but again the computational costs increase quickly. For some studies, it may be adequate to consider the paths as representing axon bundles. However, it is not yet clear if these bundles remain consistent across the volume we analyzed.

**Figure 3.**
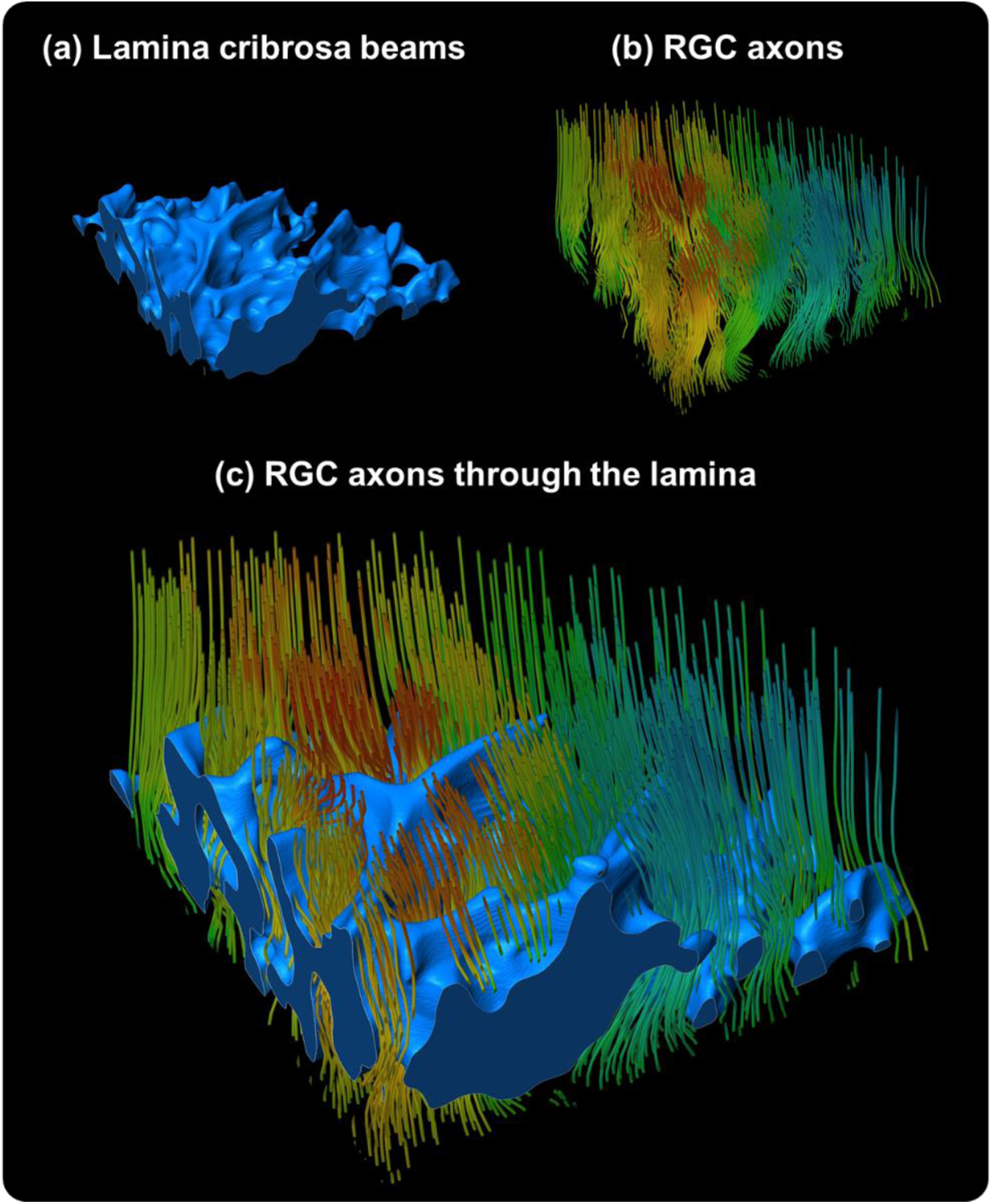
3D visualization of estimated RGC axon passing through a small region of the LC. The LC beams are shown in blue solid in (a) and (c). Estimated RGC axon paths are shown as multi-colored lines in (b) and (c), with colors indicating local axon direction. Our goal with this figure is to illustrate the tortuous axonal paths through the complex LC porous structure. Some axons seem to pass through the pores almost in a straight line, with the colors barely changing, whereas others deviate substantially, as evidenced by the color variations along their paths. The apparent discontinuities are artifacts of displaying only a small region of what is a complex 3D architecture. All the paths analyzed were continuous.

**Figure 4:**
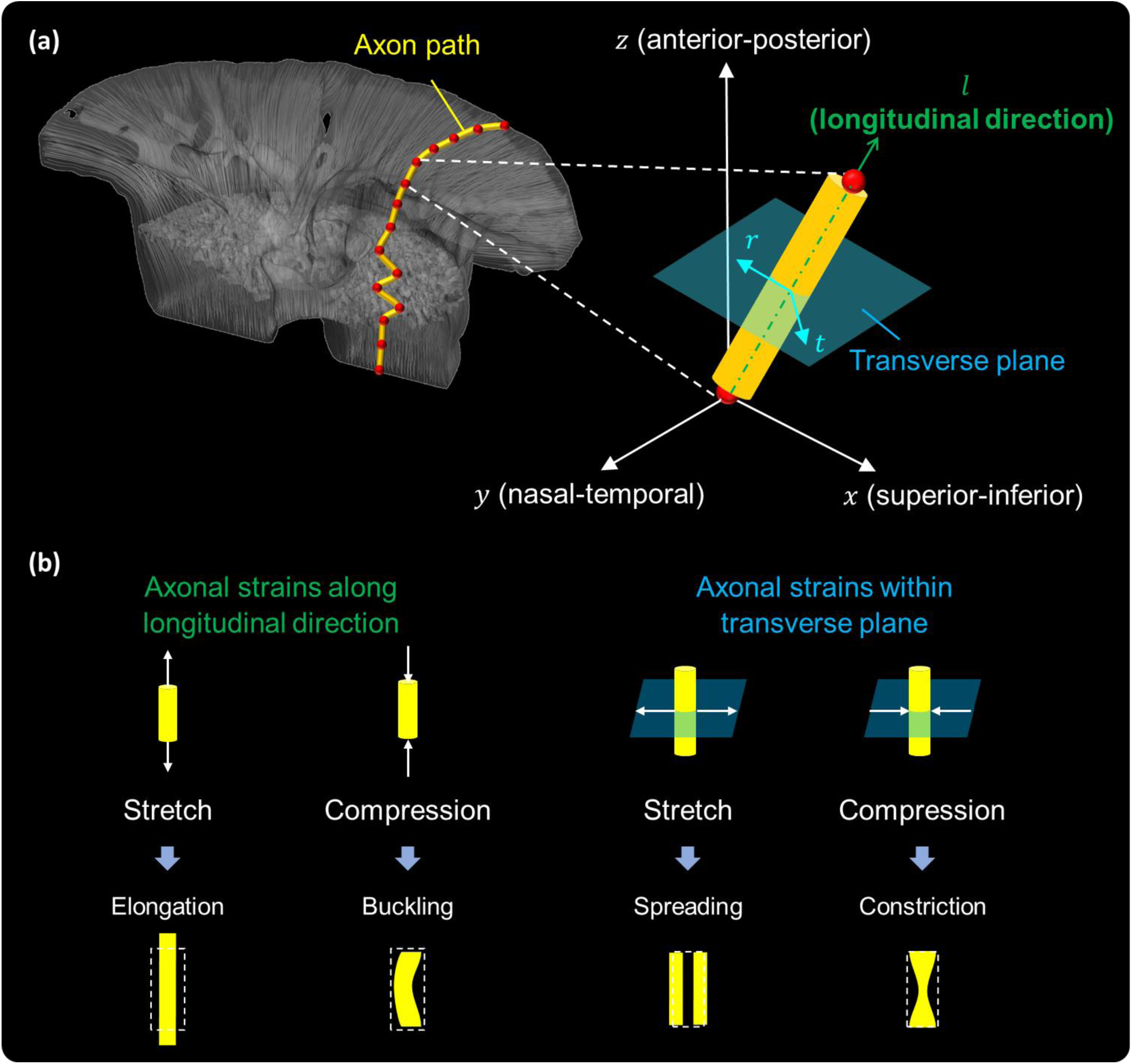
(a) Diagram illustrating a local coordinate system defined for each segment of the axon path (yellow cylinders) to quantify the insult at axon level. The axon paths, traced based on velocity from fluid mechanical simulation, consist of segments delineated by multiple axon points (red points). The local coordinate system includes a primary axis aligned longitudinally with the axon path segment (green arrow) and a transverse plane that perpendicular to the axon path segment (blue plane). This local system allows us to decompose the tissue-level strains, measured in a global Cartesian coordinate system (x-axis: superior-inferior, y-axis: nasal-temporal, z-axis: anterior-posterior), into the longitudinal and transverse axonal strains. (b) Diagram of the longitudinal and transverse axonal strains, along with their potential mechanism leading to axon damage. For instance, the longitudinal compression may cause axonal bending or buckling, while the transverse compression may induce constriction and disrupt axoplasmic transport.

The resolution and number of streamlines are directly influenced by the mesh density of the inlet surface: A denser mesh at the inlet surface results in a larger number of streamlines, while a coarser mesh results in fewer streamlines. Following too few streamlines might miss important details, while too many streamlines can significantly reduce computational efficiency and risks oversampling. Although the average monkey optic nerve has roughly 1.2 million axons, we focused on generating a sufficient number of streamlines. We deemed the number of streamlines sufficient when they capture the overall axon paths and the distribution of mechanical insults such that adding more streamlines is no longer helpful.

We want to clarify that our use of fluid mechanics to estimate axonal paths was not intended to imply that the axons are a fluid, and therefore the fluid parameters should not be interpreted as representative of fluid in the axons or similar. Our use of fluid mechanics was done based on the idea that fluid mechanics allowed us to incorporate several reasonable assumptions of axon path characteristics as detailed above (smoothness, continuity, etc.). The specific fluid properties were determined in a preliminary study to produce a smooth static laminar solution to the flow from which to obtain streamlines representative of axons. Altering the mechanical properties of the fluid or the boundary conditions driving the flow will potentially change the flow. In our preliminary tests, as long as the flow was maintained laminar the axonal paths were very similar and so were the mechanical insults on them. While we have not yet conducted a comprehensive study on the effects of the fluid properties on the flow, axon paths and results, the impression from the preliminary tests is that the effect is fairly small compared with primary factors such as the size and shape of the canal and the location of the central retinal vessels. We should also mention that fluid mechanics and closely related equations of electromagnetism have also been used successfully to describe, predict, and understand axonal paths in the retina ^31–34^. The rationale for such use ranges from the purely practical that “it works”, to more involved ones arguing that axonal paths follow potentials and solutions of diffusion of biochemical signals that ultimately translate into similar equations ^32^. In this work we lean more towards the empirical argument given the lack of sufficient information on the process by which the paths of the axons passing through the ONH are determined. Further consideration of our assumptions and discussion of the consequences are addressed in the Discussion.

### 2.5 Quantification of the axonal mechanical insult

To quantify the insult at axon level, we defined local coordinate systems for each segment of the axon path (**Figure 4**). Each system includes a primary axis aligned longitudinally with the axon path segment (*l* direction) and a transverse plane that perpendicular to it (*t* − *r* plane). The local system allows us to decompose the tissue-level strains, measured in the global Cartesian coordinate system (x-axis: superior-inferior, y-axis: nasal-temporal, z-axis: anterior-posterior), into the longitudinal and transverse axonal strains. The transformation is achieved via a rotation matrix ***Q***:

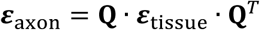

where ***ε***_axon_ and ***ε***_tissue_ represent the mechanical insult at axon and tissue levels, respectively. At each point along the axon path, ***ε***_axon_ includes a normal strain component in the longitudinal direction (*ε*_*ll*_), and normal and shear strain components within the transverse plane. The transverse axonal insult is determined by the axon cross-sectional area change using two principal strains, 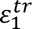 and 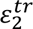:

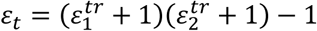

A positive *ε*_*t*_ denotes transverse stretch (cross-section area increase), while a negative *ε*_*t*_indicates transverse compression (cross-section area decrease).

#### A note on terminology

The term “axonal strain” in our paper refers to components of tissue-level strains resolved into longitudinal and transverse directions relative to the axon paths. Due to the limitations of OCT image resolution and quality, these are not actual axon-level strains. We use the term “tissue-level strain” to represent the conventional calculations of strain, including stretch, compression, shear, and effective strains.

### 2.6 Analysis of axonal insults

To evaluate potential advantages of the axon-centric approach, we analyzed the axonal insults through four detailed examples.

#### Comparison of strain distributions at tissue and axon levels in ONH

For both strains, we colored the estimated axon paths with the strain values. A traditional physical color bar was used to represent tissue-level strain, with red indicating high strain and blue indicating low. A new color bar was introduced for axonal strain: white to blue to cyan for increasing stretch (positive strain values), and white to red to yellow indicating increasing compression (negative strain values). Additionally, strain distributions were shown on an axial section of mechanical insults at tissue and axon levels.

#### Axonal insults along two estimated axon paths

We showed the axonal strain changes along individual axons, connecting the RNFL to the optic nerve. We also calculated the maximum and cumulative axonal insult values for both paths.

#### Volumetric distribution of axonal insults

We assumed uniform circular cross-sections for all axon paths. All points from these paths were used to construct the histogram plots, illustrating the proportion of total axonal volume undergoing each level of strain. Bin heights in the histogram represented the percentage of axonal volume under each strain level, with bins set at fixed 0.2% strain increments.

#### Spatial distribution of axonal insults

Spatial distribution plots of both longitudinal and transverse axonal strain were created to illustrate the spatial density of axonal insults across the ONH regions from anterior (RNFL) to posterior (optic nerve). Scatter points were plotted based on its axonal strain value and location in the ONH. We applied kernel density estimation, a nonparametric technique, to estimate the density of these scatter points. The resulting spatial distribution was visually represented through a color gradient, with high-density areas depicted in red and low-density areas in purple.

## 3. Results

The distribution of mechanical insults at both tissue and axon levels within ONH is shown in **Figure 5**. For the strains at tissue level, the axons undergo non-uniform compression and stretch, while their directions are unspecified. No clear patterns were identified for tissue-level strains. However, for the axonal strains, a notable pattern was observed: axons in the LC and post-LC areas predominantly experienced longitudinal compression and transverse stretch, whereas those in the pre-LC and ONH rim mainly suffered longitudinal stretch and transverse compression.

**Figure 5.**
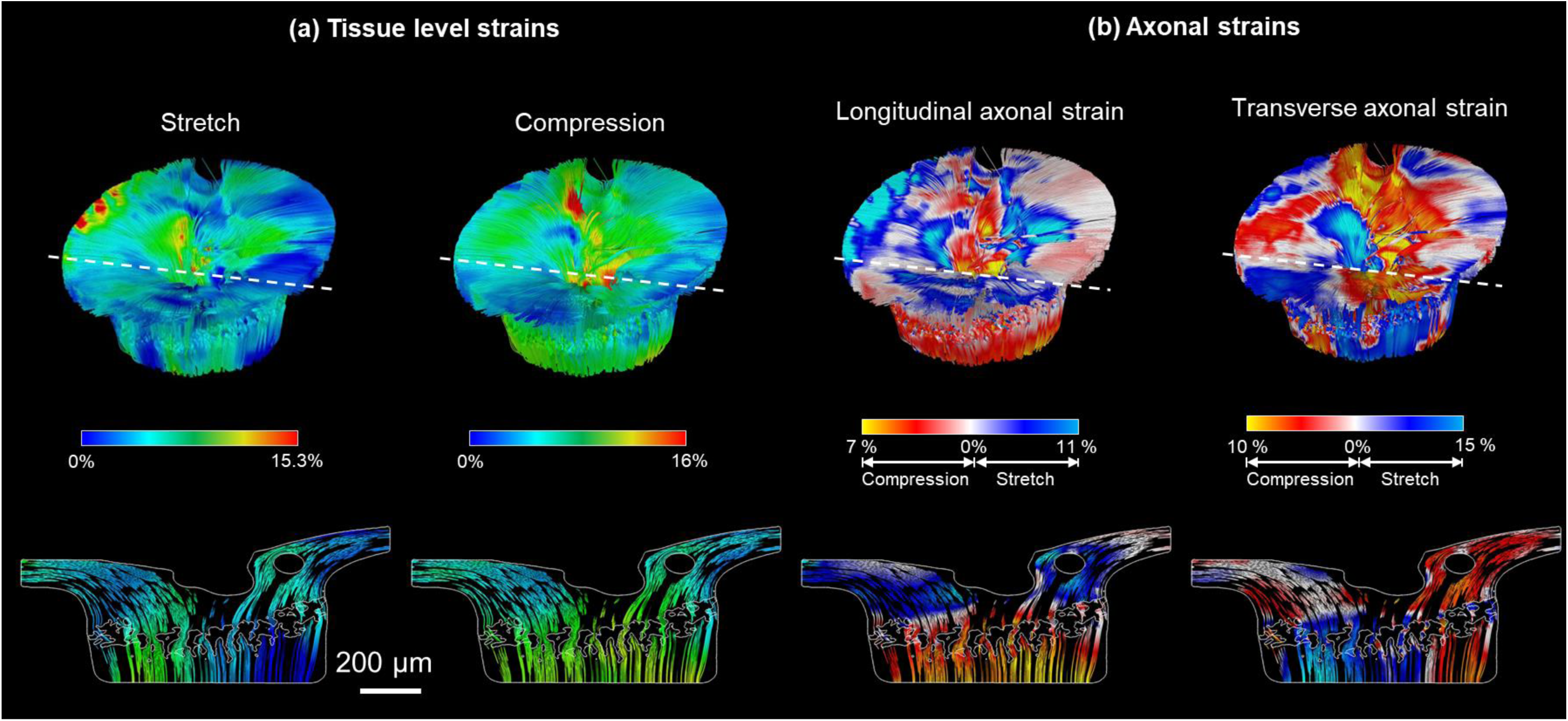
Strain distribution in ONH at the tissue and axon levels with the strain values colored on the estimated axon paths: (a) strain at tissue level, where red highlights areas of high strain and blue indicates low. (b) strain at the axon level, with a different color bar: white to red to yellow signifies increasing levels of compression (negative strain values), while white to blue to cyan represents increasing levels of stretch (positive strain values). The bottom row of the figure displays axial sections of those strains, with the location of the section indicated by a white dashed line on the 3D distribution (top row). For the strains at tissue level, the axons undergo non-uniform compression and stretch, while their directions are unspecified. For the strains at axon level, a distinct patten is observed: axons in the LC and post-LC areas predominantly experience longitudinal compression and transverse stretch, whereas those in the pre-LC and retina rim mainly undergo longitudinal stretch and transverse compression.

Distribution and cumulative effect of the strains at axon level (longitudinal and transverse) along two representative axons is shown in **Figure 6**. From posterior to anterior along both axons, the longitudinal axonal strain changes from compression to stretch, while the transverse axonal strain changes from stretch to compression. For transverse compression, the maximum value for axon 1 is 33% lower compared to axon 2, while the cumulative value for axon 1 is 30% higher than that in axon 2. This indicates that some axons, even with lower local maximum insult values, might suffer injury due to high cumulative insult. There was no clear overall relationship observed between the maximum and cumulative insult along individual axons.

**Figure 6.**
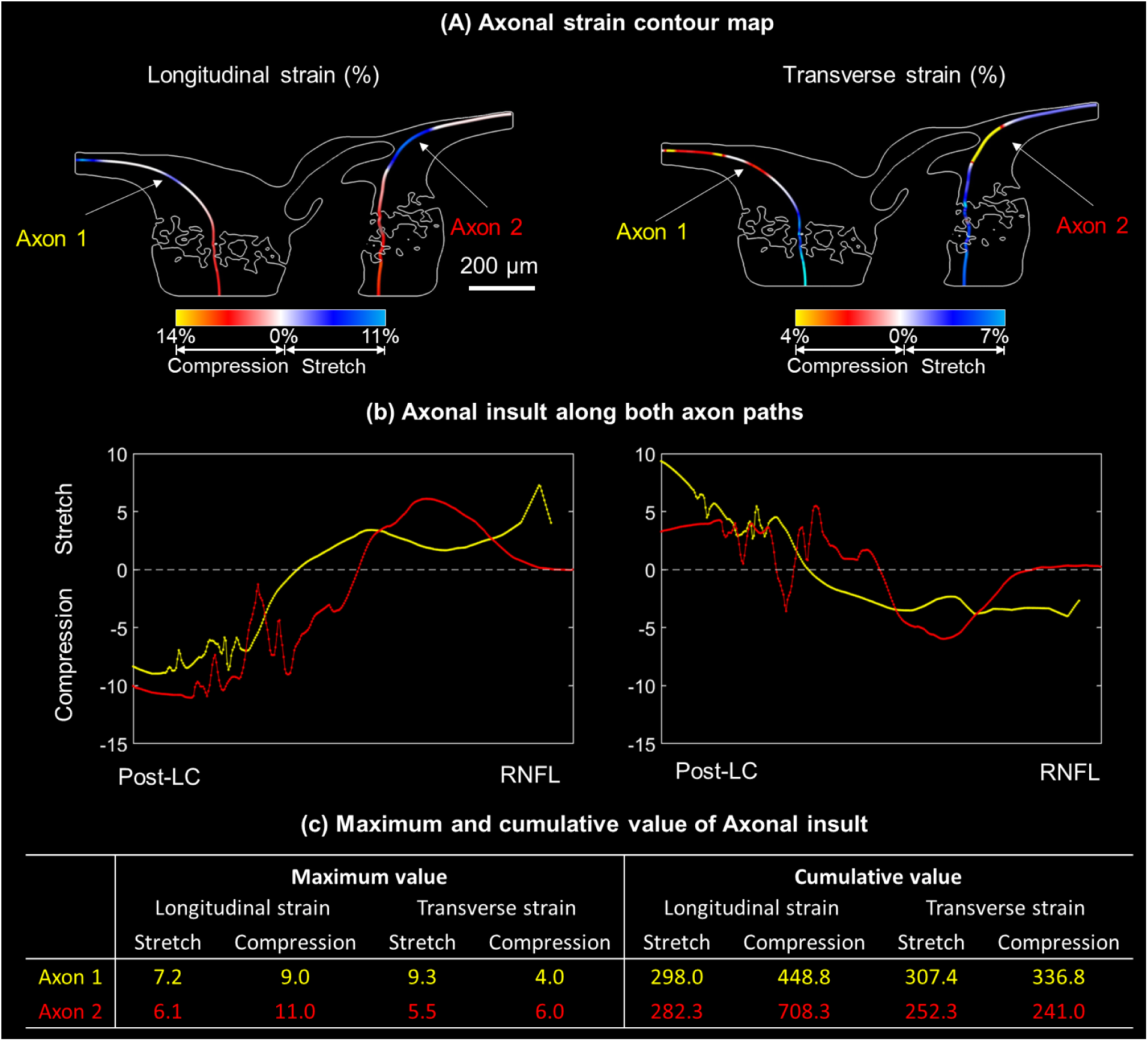
The longitudinal (left) and transverse (right) axonal strain along two example axons. Axon 1 (data shown in yellow) is from the temporal region and axon 2 (data shown in red) is from the nasal region. **(a)** Contour map of the strains along the axon. Axons were colored according to the magnitude of stretch (white to cyan) and compression (white to yellow). **(b)** The magnitude of both axonal strains for points along the length of axon from posterior to anterior. From posterior to anterior along both axons, the longitudinal axonal strain changes from compression to stretch, while the transverse axonal strain changes from stretch to compression. **(c)** The maximum and cumulative value of the axonal insults for both axons. For transverse compression, the maximum value for axon 1 is 33% lower compared to axon 2, while the cumulative value for axon 1 is 30% higher than that in axon 2.

Volumetric distribution plots of the longitudinal and the transverse axonal strain are shown in **Figure 7**, illustrating the proportion of axonal volume undergoing each level of axonal strain. For longitudinal strain, 50.3% of the axonal volume is stretched, and 49.7% is compressed. In contrast, for transverse strain, 31.8% is stretched and 68.2% is compressed.

**Figure 7.**
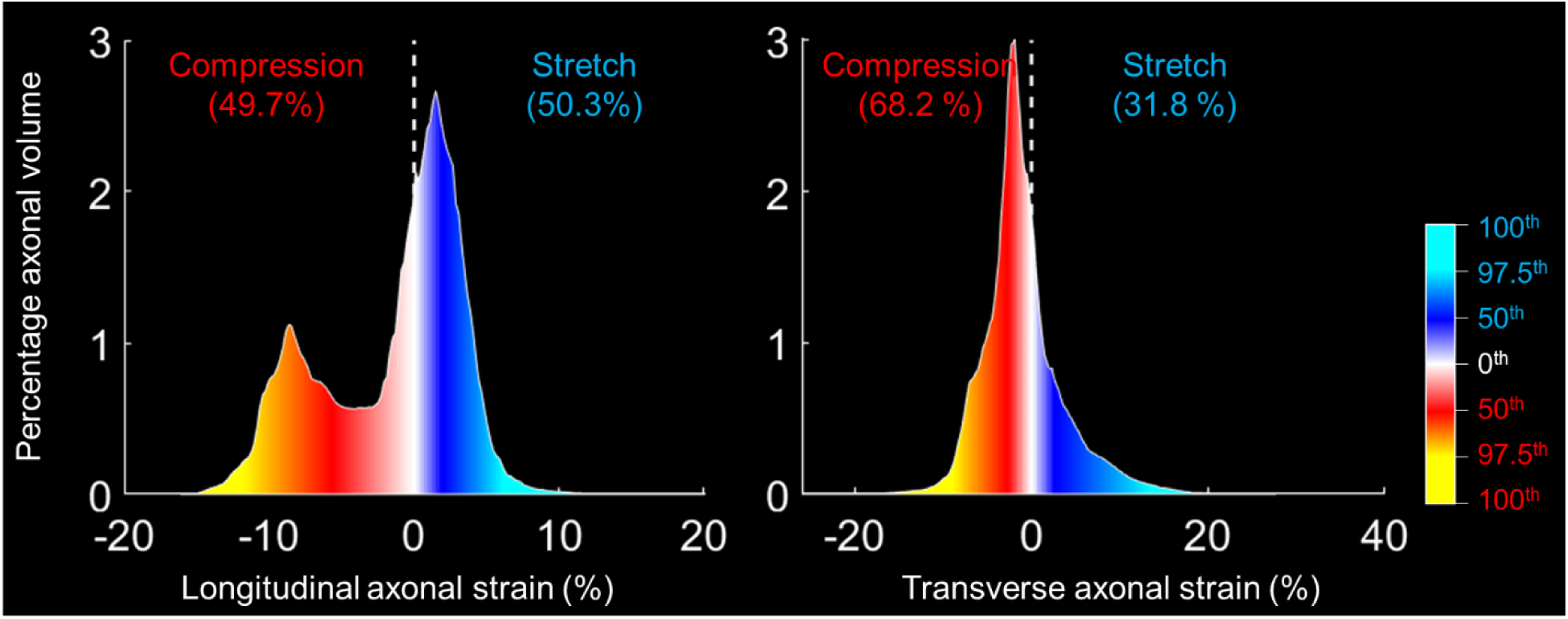
Volumetric distribution plot of the longitudinal axonal strain (left) and the transverse axonal strain (right). The color spectrum indicates the degree of strain: white to blue to cyan represents increasing levels of stretch (positive strain values), while white to red to yellow signifies increasing levels of compression (negative strain values). All points from these paths were used to construct the histogram plots, illustrating the proportion of axonal volume undergoing each level of strain. The bins have a fixed width representing 0.2% strain increments. For longitudinal strain, 50.3% of the axonal volume is stretched, and 49.7% is compressed. In contrast, for transverse strain, 31.8% is stretched and 68.2% is compressed.

The spatial distributions of the mechanical insults at axon level are shown in **Figure 8**, illustrating the spatial density of axonal insults across the ONH regions from anterior (RNFL) to posterior (optic nerve). For example, the highest density of longitudinal stretch was observed for the portion of axons within the RNFL and ONH rim.

**Figure 8.**
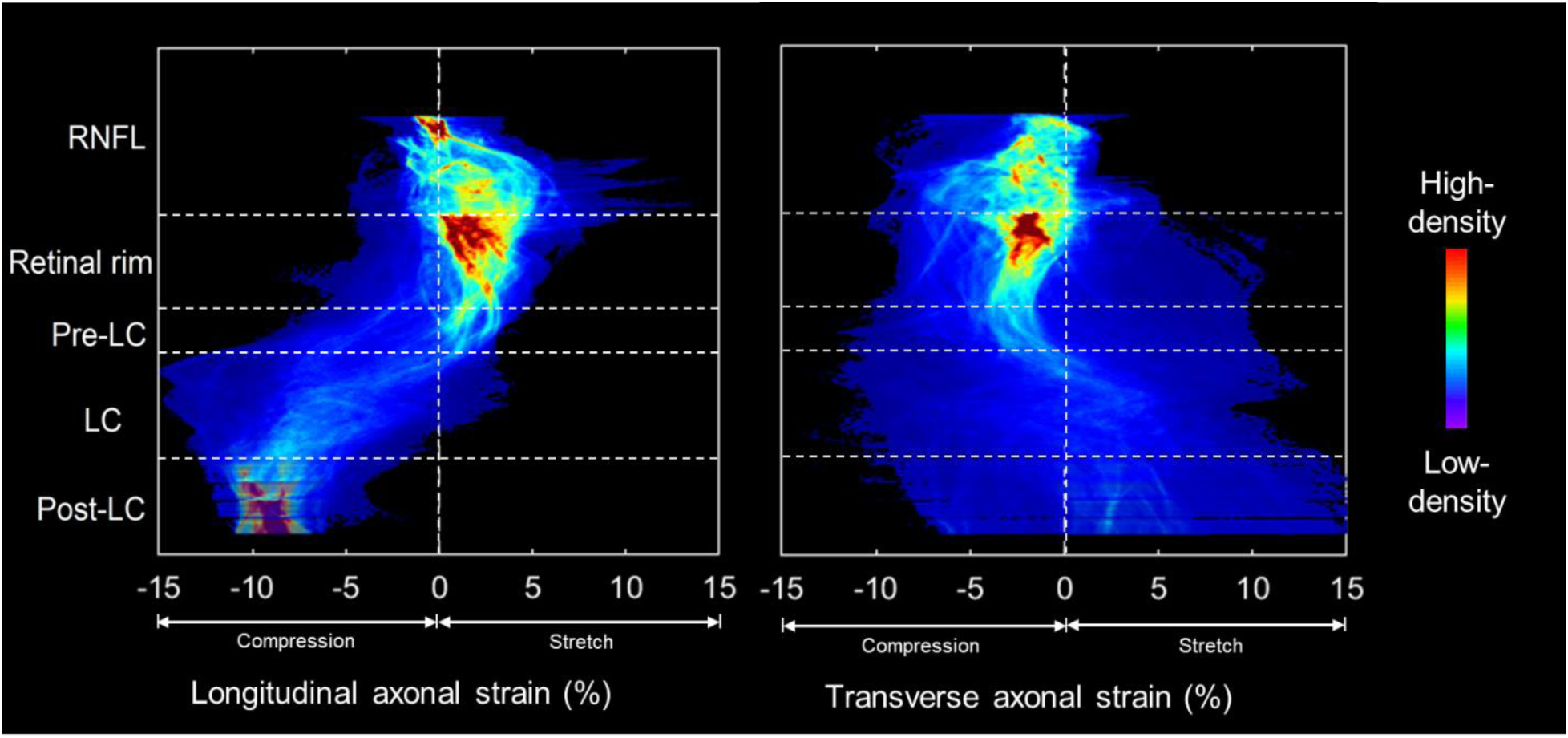
Spatial distribution plots of longitudinal (left) and transverse (right) axonal strain, illustrating the spatial density of the insult across the ONH region from the anterior (RNFL) to the posterior (optic nerve). The kernel density function was employed to determine the density of scatter points, each representing an axonal point plotted with its axonal strain value as the x-coordinate and its ONH location as the y-coordinate. The spatial distribution was visually represented through a color gradient, with high-density areas depicted in red and low-density areas in purple. For example, the highest density of longitudinal stretch was observed for the portion of axons within the RNFL and ONH rim.

## 4. Discussion

Our goal was to describe and demonstrate an axon-centric approach for quantifying the mechanical insult to RGC axons within the ONH under elevated IOP. Specifically, we utilized experimental ONH deformation data, along with approximation of continuous 3D axon paths, to quantify IOP-induced insult at axon level based on their local path orientation. We analyzed the IOP-induced ONH deformation in a monkey eye to illustrate the potential of the axon-centric approach.

We highlight two main advantages of the axon-centric approach over earlier approaches. First, it allows estimation of axonal mechanical insults longitudinally and transversely. Second, it allows visualization of the insult along individual axon path, providing a novel link to relate regions from RNFL to the LC via individual axons. Below we discuss these advantages in detail, along with the motivation and the rationale for our study, followed by an extensive discussion of the assumptions, limitations and other considerations to keep in mind when interpreting the work.

*Advantage 1: the axon-centric approach allows estimation of axonal mechanical insults longitudinally and transversely*.

The axon-centric approach allows for quantifying axonal insults within the ONH in both longitudinal and transverse directions. IOP-induced mechanical insult on retinal ganglion cell axons passing through the optic nerve head (ONH) is believed to be a major factor in axonal damage and glaucoma ^3, 4, 6, 7^. However, current experimental and computational methods primarily focus on measuring or calculating tissue-level mechanical insult, overlooking the distinct spatial characteristics of individual axons. The axon-centric approach enables a detailed analysis of the mechanical insults at axon level, considering their unique geometric properties and susceptibilities.

Axonal mechanical insult involves multidirectional stretch and compression, potentially leading to axonal damage through different mechanisms. For instance, compression transverse to an axon could adversely affect axonal transport and/or axoplasmic flow, whereas compression longitudinally to an axon may cause structural instabilities such as axonal buckling or deformations that trigger different types of mechanoreceptors ^22, 23, 35–38^. The influence of the magnitude of local stretch and compression on axonal damage is incompletely understood. It is plausible that a lower magnitude of transverse compression may induce more axonal damage compared to a higher magnitude of transverse stretch. Therefore, even minor transverse compressive insults should not be overlooked as potential contributors to axonal injury. This information is valuable to establish a clearer connection between IOP-induced insult and glaucoma. It should be noted that the compression along the axonal direction can potentially induce buckling due to the high aspect ratio of axons. Buckling is a highly unstable structure response that may cause axons to change their tortuosity or angles at specific locations. It seems plausible that this could disrupt axonal functions, for instance by disrupting mitochondrial transport, but to the best of our knowledge, experimental evidence of these effects is not yet available.

*Advantage 2: the axon-centric approach allows visualization of the insult along individual axon path, providing a novel link to relate regions from RNFL to the LC via individual axons*.

In conventional regional studies, the LC and RNFL are divided into the temporal, superior, nasal, and inferior regions ^39, 40^. It should be noted that axon paths from the RNFL to the optic nerve do not always stay within a single quadrant; they may navigate around LC beams or large blood vessels, occasionally bending or twisting, and even crossing from the temporal directly into the nasal region. Our estimated axon paths, spanning from the RNFL to the optic nerve, offer a novel link to relate these regions. Furthermore, our approach allows for characterizing both local maximum and cumulative insult along the length of individual axons.

For instance, as observed in **Figure 6**, along the axon, longitudinal strain transitions from stretch to compression, and transverse strain shifts from compression to stretch as the axon enters the LC region. In transverse compression, axon 1 exhibits a maximum value that is 33% lower than axon 2, yet its cumulative value is 30% higher. Those results suggest that axons with lower maximum insults might sustain injury due to high cumulative insults. Conversely, axons with similar cumulative insults could face different damage risks if the local maximum insult is higher in one than in another. This comprehensive analysis not only enhances our understanding of axonal behavior within the LC and RNFL regions but also aids in establishing the connection between IOP-induced axonal insult and the development of glaucoma.

We are not aware of any previous reports that have quantified IOP-induced mechanical insult within the ONH with an axon-centric approach. Doing this required information about axon orientations and paths from the RNFL, through the ONH, including the LC, and to the distal optic nerve. Although techniques for mapping axonal paths within the RNFL and to the edges of the disc have been demonstrated with excellent results even in-vivo ^41–44^ (with some of the latest approaches taking advantage of polarization-sensitive OCT for high-resolution mapping ^45^), the techniques do not extend to large ONHs with collagenous LCs. Axonal tracing through the rodent ONH has been possible ^46^ due to the smaller size, and the lack of collagen in their glial lamina. Collagenous components in monkey and human lamina make axonal paths more complex and make imaging difficult through scattering and absorption. Whilst labeling neural tissue components like axons and astrocytes in the ONH is feasible ^47–50^, differentiating individual axons or bundles for tracing remains challenging.

A few techniques have been deployed for visualizing and tracing a few axons through the ONH, often selected semi-randomly or based on a common origin in the RNFL, optic nerve or brain ^51–58^. A fine example of this approach is the work by Morgan and colleagues studying the course of RGC axons through the LC using horseradish peroxidase tracer applied at the optic nerve ^59^. They found that the “majority of axons took a direct course through the lamina cribrosa but a significant minority, in the 8-12%, deviated to pass between the cribrosal plates in both central and peripheral parts of the optic disc.” Although we did not quantify the fraction of axons that pass directly through the LC or not, our analysis allowed for axons to pass between beams, some exhibiting paths that would appear as the deviations in Morgan et al.’s paper (**Figure 2**). Morgan et al. postulated that the deviated axons may be more vulnerable to compression. This is consistent with our interpretations as the “deviated” axons would be subject to different insult.

Leveraging axonal transport other studies have done spatial mappings between the retina and optic nerve ^43^. The studies, however, did not report axonal paths through the LC. Several techniques have been reported to quantitatively analyze axonal health at the optic nerve ^60–63^. These include tools for map axons within the optic nerve, as these are used routinely to study axonal damage, for example by crush, and/or regeneration ^64^. To the best of our knowledge no axonal path data through the collagenous LC was reported.

As noted in the methods, for the demonstration model used in this manuscript the detailed LC beam reconstruction and the OCT delineations – deep and shallow tissues, respectively, were obtained from contralateral eyes. Both reconstructions were made of healthy tissues from the same animal, and are therefore consistent with one another, but they were not from the same eye. Elsewhere, several studies have demonstrated that contralateral eyes, when healthy, as in this case, are substantially and significantly more similar than unrelated eyes ^13, 65–70^. Importantly, this applies to larger scale morphology ^66–69, 71^, meso-scale characteristics of the beams and pores ^13, 69, 70, 72, 73^, and even to micro-scale collagen fiber crimp^74–76^. It was only after we had completed the reconstructions that we realized that the eye we had selected for OCT reconstruction had later been the one used to study unilateral longitudinal experimental glaucoma. Because of this, the histology of this eye available after sacrifice no longer represented a healthy eye. Thus, while we acknowledge that, ideally, we would have used data only from one eye, we posit that the effects of mixing data from contralateral eyes are limited and thus that this work is still useful as a proof-of-principle. Future work should evaluate the consequences of this decision.

In our study, we used the experimental deformations from the DVC of OCT together with the axonal orientations to obtain the axonal strains. However, numerical studies highlight the mismatch in elastic properties between neural tissues and collagen beams results in a larger strain in the neural tissue ^77–79^. Like the work we present in this manuscript, computational modeling studies contain results that are reasonable but that have yet to be fully verified experimentally. This work does not dispute those results. Instead, we present an alternative methodology to quantify the various forms of insult to the axons. Our new method is rooted in the experimental information from DVC of OCT together with a detailed LC reconstruction and axonal path prediction. By being based on DVC-derived experimental data, the work on this manuscript does not require some potentially crucial assumptions made in computational modeling, such as the perfect attachment between neural and connective tissues, the tissues incompressibility, the constitutive formulations for each component, etc. If these assumptions do not hold, the predictions from the models might change substantially. Some aspects of the new method we propose, like transforming strains from the tissue-level perspective into axonal perspective strains can be learned from this paper and applied also to computational models. We welcome this possibility and would argue that considering the orientation of the axons in their constitutive properties will potentially help improve the numerical predictions.

It is important to acknowledge other limitations of this work. First, all the observations reported herein were obtained from a single healthy female monkey eye. Thus, conclusions about population-level findings cannot be drawn until eyes from more animals are analyzed. Future studies will aim to analyze multiple animals in both sexes to provide a more comprehensive understanding of the IOP-induced axonal effects. Moreover, it is possible that glaucomatous eyes exhibit different deformations with acutely elevated IOP compared to healthy eyes. Future work will involve analyzing a larger number of healthy eyes, and applying this analysis to glaucomatous eyes, in order to advance our understanding of IOP-induced axonal insults within the ONH.

Second, the axonal paths were estimated as the streamlines of a fluid. Axons are not fluid, and their paths are not required to follow streamlines. The rationale for this choice was presented in the Methods section and the **Supplementary document S1**. Briefly, it allowed us to incorporate several reasonable assumptions about the axons and their paths. Further, as noted above, similar methods have been proven successful for estimating axonal paths on the RNFL ^41–43^. We acknowledge that the axon-centric approach may be imperfect, but we still posit that it provides very reasonable first approximations to the 3D axon paths across the ONH. The directional information of the estimated axonal paths is crucial to transform tissue-level insult to axon-type insult. While the techniques for measuring axonal paths precisely within the collagenous LC remain out of reach, several current techniques allow visualization of the more superficial tissues of the ONH. An example is shown in the new **Supplementary Figure 4**. Future work should evaluate if axonal paths must be determined even more accurately, or perhaps that a simpler approach could be sufficient. For instance, it may be enough to assume that the axons are along their primary anterior-posterior direction. The approach we describe is a reasonable starting point.

Third, the 3D LC beam structure used in this study was sourced from histological images of a different monkey eye. This is because there are vascular shadows OCT images in the LC region due to technical limitations. LC beam structure derived from OCT scans could miss important structure and impede the estimation of axon paths. As an alternative, the reconstruction of LC beams was based on histological images of a different monkey eye. While IPOL images provide clear details of LC structure, it is important to note that ONH morphology, especially LC beam structure, can vary between eyes. This variability may influence axon path estimation and, subsequently, the quantification of axonal insults. Future studies will include LC beam structures from the same eyes as those imaged by OCT, as well as evaluate the effect of differing LC structure on estimates of axonal insults to address this question. Moreover, clinical applications of this approach will require that all the predictions are made based on in vivo data, without any need for histology.

Fourth, it is possible that the tissues distort during cryosectioning and mounting. Elsewhere we have shown that these deformations are small and LC structures are comparable between OCT and histology (Wang et al. 2017). Nevertheless, any distortions will affect the accuracy of the axon paths calculated.

Fifth, in the reconstruction of non-collagenous axonal volume, we did not consider other elements of the LC and ONH, such as capillaries or astrocytes. There are several major challenges to including these microscale features, in large part due to their enormous complexity for both the vascular network ^26, 80, 81^ and astrocytes ^52, 82^. Adding these features will therefore represent a huge increase in the challenges for reconstruction and computational cost. An even larger challenge is that neither of these components can be visualized or mapped in vivo. On the positive side, it remains unclear whether it is crucial to consider these elements to make useful predictions of axonal paths. As discussed before, useful maps of axonal paths within the RNFL have been made. To the best of our knowledge, none of them incorporated smaller scale features of microvasculature or glia.

Sixth, due to the current resolution limitations of OCT imaging, the DVC analysis represents some degree of homogenization. In regions like the LC incorporating components with very distinct mechanics, such as neural tissues and collagen, the homogenization may not accurately capture the true axonal strains. In other regions, like the rim, the pre-LC and retro-LC, with tissues that likely differ less in mechanical properties, like axons and astrocytes, the DVC-calculated deformations are potentially more accurate. We recognize that axons may deform independently, potentially leading to complex deformations, which could disrupt axon-astrocyte bonds and lead to remodeling, inflammation, or apoptosis. Developing tools to measure neural tissue deformations at a smaller scale is crucial. However, current techniques for visualizing axons within the collagenous ONH are limited, making it challenging to directly test their deformations due to elevated IOP. For this work, we argue that it is reasonable to use as a first approximation the assumption that axonal deformations are aligned with the measurements obtained from the measurements obtained through DVC.

Seventh, we have not carried out a comprehensive sensitivity study on model parameters. These represent an important area of future research. We note that the predictions did not substantially change with the fluid mechanics properties, suggesting that the paths predicted with this technique are potentially useful.

*<u>In summary</u>*, we posit that the proposed axon-centric approach is useful as a first approximation to quantify the IOP-induced mechanical insult to RGC axons within the ONH. Our approach distinguishes the mechanical insult on axons longitudinally and transversely, suggesting distinct mechanisms of potential axon damage. It enabled the estimation of axon paths based on the in vivo data before the axon is damaged. The results can be later used to relate the IOP-induced axonal insults with the chronic neural tissue loss. This approach can potentially help understand the biomechanics of ONH and its relationship to glaucoma.

## Acknowledgements

Supported in part by Research to Prevent Blindness, National Institutes of Health R01-EY023966, R01-EY030590, R01-EY031708, T32-EY017271, P30-EY008098, Canadian Institutes of Health Research PJT-178008, Eye and Ear Foundation (Pittsburgh, PA), Research to Prevent Blindness (unrestricted grant to UPMC’s Department of Ophthalmology and Stein Innovation Award to IA Sigal).

## References

1. Quigley HA, Addicks EM, Green WR, Maumenee AE. Optic Nerve Damage in Human Glaucoma: II. The Site of Injury and Susceptibility to Damage. Archives of Ophthalmology 1981;99:635–649.

2. Gaasterland D, Tanishima T, Kuwabara T. Axoplasmic flow during chronic experimental glaucoma. 1. Light and electron microscopic studies of the monkey optic nervehead during development of glaucomatous cupping. Investigative Ophthalmology & Visual Science 1978;17:838–846.

3. Quigley HA, Hohman RM, Addicks EM, Massof RW, Green WR. Morphologic Changes in the Lamina Cribrosa Correlated with Neural Loss in Open-Angle Glaucoma. American Journal of Ophthalmology 1983;95:673–691.

4. Quigley HA. Glaucoma: macrocosm to microcosm the Friedenwald lecture. Investigative Ophthalmology & Visual Science 2005;46:2662–2670.

5. Heijl A, Leske MC, Bengtsson B, et al. Reduction of intraocular pressure and glaucoma progression: results from the Early Manifest Glaucoma Trial. Archives of ophthalmology 2002;120:1268–1279.

6. Sigal IA, Ethier CR. Biomechanics of the optic nerve head. Experimental Eye Research 2009;88:799–807.

7. Gardiner SK, Fortune B, Wang L, Downs JC, Burgoyne CF. Intraocular pressure magnitude and variability as predictors of rates of structural change in non-human primate experimental glaucoma. Exp Eye Res 2012;103:1–8.

8. Singh R, Zhao Y, Elze T, et al. Polygenic Risk Score Improves Prediction of Primary Open Angle Glaucoma Onset in the Ocular Hypertension Treatment Study. medRxiv 2023;2023.2008.2015.23294141.

9. Van Gelder RN, Chiang MF, Dyer MA, et al. Regenerative and restorative medicine for eye disease. Nature Medicine 2022;28:1149–1156.

10. Stowell C, Burgoyne CF, Tamm ER, Ethier CR, Lasker IIoA, Glaucomatous Neurodegeneration P. Biomechanical aspects of axonal damage in glaucoma: A brief review. Exp Eye Res 2017;157:13–19.

11. Girard MJ, Strouthidis NG, Desjardins A, Mari JM, Ethier CR. In vivo optic nerve head biomechanics: performance testing of a three-dimensional tracking algorithm. Journal of The Royal Society Interface 2013;10:20130459.

12. Sigal IA, Grimm JL, Jan N-J, Reid K, Minckler DS, Brown DJ. Eye-Specific IOP-Induced Displacements and Deformations of Human Lamina Cribrosa. Investigative Ophthalmology & Visual Science 2014;55:1–15.

13. Coudrillier B, Campbell IC, Read AT, et al. Effects of Peripapillary Scleral Stiffening on the Deformation of the Lamina Cribrosa. Investigative Ophthalmology & Visual Science 2016;57:2666–2677.

14. Fazio MA, Clark ME, Bruno L, Girkin CA. In vivo optic nerve head mechanical response to intraocular and cerebrospinal fluid pressure: imaging protocol and quantification method. Scientific Reports 2018;8:12639.

15. Behkam R, Kollech HG, Jana A, et al. Racioethnic differences in the biomechanical response of the lamina cribrosa. Acta Biomaterialia 2019;88:131–140.

16. Bellezza AJ, Hart RT, Burgoyne CF. The Optic Nerve Head as a Biomechanical Structure: Initial Finite Element Modeling. Investigative Ophthalmology & Visual Science 2000;41:2991–3000.

17. Sigal IA, Flanagan JG, Tertinegg I, Ethier CR. Finite Element Modeling of Optic Nerve Head Biomechanics. Investigative Ophthalmology & Visual Science 2004;45:4378–4387.

18. Roberts MD, Liang Y, Sigal IA, et al. Correlation between Local Stress and Strain and Lamina Cribrosa Connective Tissue Volume Fraction in Normal Monkey Eyes. Investigative Ophthalmology & Visual Science 2010;51:295–307.

19. Fazio MA, Grytz R, Bruno L, et al. Regional Variations in Mechanical Strain in the Posterior Human Sclera. Investigative Ophthalmology & Visual Science 2012;53:5326–5333.

20. Weinreb RN, Leung CKS, Crowston JG, et al. Primary open-angle glaucoma. Nature Reviews Disease Primers 2016;2:16067.

21. Dias MS, Luo X, Ribas VT, Petrs-Silva H, Koch JC. The Role of Axonal Transport in Glaucoma. International Journal of Molecular Sciences 2022;23:3935.

22. Yu D-Y, Cringle SJ, Balaratnasingam C, Morgan WH, Yu PK, Su E-N. Retinal ganglion cells: Energetics, compartmentation, axonal transport, cytoskeletons and vulnerability. Progress in Retinal and Eye Research 2013;36:217–246.

23. Anderson DR, Hendrickson A. Effect of intraocular pressure on rapid axoplasmic transport in monkey optic nerve. Invest Ophthalmol 1974;13:771–783.

24. Tamm ER, Ethier CR, Lasker IIoA, Glaucomatous Neurodegeneration P. Biological aspects of axonal damage in glaucoma: A brief review. Exp Eye Res 2017;157:5–12.

25. Zhong F, Wang B, Wei J, et al. A high-accuracy and high-efficiency digital volume correlation method to characterize in-vivo optic nerve head biomechanics from optical coherence tomography. Acta Biomaterialia 2022.

26. Waxman S, Brazile BL, Yang B, et al. Lamina cribrosa vessel and collagen beam networks are distinct. Experimental Eye Research 2022;215:108916.

27. Yang B, Lee P-Y, Hua Y, et al. Instant polarized light microscopy for imaging collagen microarchitecture and dynamics. Journal of Biophotonics 2021;14:e202000326.

28. Tran H, Wallace J, Zhu Z, et al. Seeing the Hidden Lamina: Effects of Exsanguination on the Optic Nerve Head. Investigative Ophthalmology & Visual Science 2018;59:2564–2575.

29. Wang B, Tran H, Smith MA, et al. In-vivo effects of intraocular and intracranial pressures on the lamina cribrosa microstructure. PLoS One 2017;12:e0188302.

30. Tran H, Grimm J, Wang B, et al. Mapping in-vivo optic nerve head strains caused by intraocular and intracranial pressures. Proc SPIE Int Soc Opt Eng 2017;10067.

31. Tan O, Liu L, Liu L, Huang D. Nerve Fiber Flux Analysis Using Wide-Field Swept-Source Optical Coherence Tomography. Translational Vision Science & Technology 2018;7:16–16.

32. Airaksinen PJ, Doro S, Veijola J. Conformal geometry of the retinal nerve fiber layer. Proceedings of the National Academy of Sciences 2008;105:19690–19695.

33. Jansonius NM, Schiefer J, Nevalainen J, Paetzold J, Schiefer U. A mathematical model for describing the retinal nerve fiber bundle trajectories in the human eye: Average course, variability, and influence of refraction, optic disc size and optic disc position. Experimental Eye Research 2012;105:70–78.

34. Jansonius NM, Nevalainen J, Selig B, et al. A mathematical description of nerve fiber bundle trajectories and their variability in the human retina. Vision Research 2009;49:2157–2163.

35. Vicic N, Guo X, Chan D, Flanagan JG, Sigal IA, Sivak JM. Evidence of an Annexin A4 mediated plasma membrane repair response to biomechanical strain associated with glaucoma pathogenesis. Journal of Cellular Physiology 2022;237:3687–3702.

36. Alqawlaq S, Flanagan JG, Sivak JM. All roads lead to glaucoma: Induced retinal injury cascades contribute to a common neurodegenerative outcome. Experimental Eye Research 2019;183:88–97.

37. Morrison JC. Integrins in the optic nerve head: potential roles in glaucomatous optic neuropathy (an American Ophthalmological Society thesis). Transactions of the American Ophthalmological Society 2006;104:453.

38. Irnaten M, O’Brien CJ. Calcium-Signalling in Human Glaucoma Lamina Cribrosa Myofibroblasts. International Journal of Molecular Sciences 2023;24:1287.

39. He L, Yang H, Gardiner SK, et al. Longitudinal Detection of Optic Nerve Head Changes by Spectral Domain Optical Coherence Tomography in Early Experimental Glaucoma. Investigative Ophthalmology & Visual Science 2014;55:574–586.

40. Midgett D, Liu B, Ling YTT, Jefferys JL, Quigley HA, Nguyen TD. The Effects of Glaucoma on the Pressure-Induced Strain Response of the Human Lamina Cribrosa. Investigative Ophthalmology & Visual Science 2020;61:41–41.

41. Grannonico M, Miller DA, Liu M, et al. Global and Regional Damages in Retinal Ganglion Cell Axon Bundles Monitored Non-Invasively by Visible-Light Optical Coherence Tomography Fibergraphy. J Neurosci 2021;41:10179–10193.

42. Miller DA, Grannonico M, Liu M, et al. Visible-Light Optical Coherence Tomography Fibergraphy for Quantitative Imaging of Retinal Ganglion Cell Axon Bundles. Transl Vis Sci Technol 2020;9:11.

43. Naito J. Retinogeniculate projection fibers in the monkey optic nerve: a demonstration of the fiber pathways by retrograde axonal transport of WGA-HRP. J Comp Neurol 1989;284:174–186.

44. Carreras FJ, Medina J, Ruiz-Lozano M, Carreras I, Castro JL. Virtual Tissue Engineering and Optic Pathways: Plotting the Course of the Axons in the Retinal Nerve Fiber Layer. Investigative Ophthalmology & Visual Science 2014;55:3107–3119.

45. Sugita M, Pircher M, Zotter S, et al. Retinal nerve fiber bundle tracing and analysis in human eye by polarization sensitive OCT. Biomed Opt Express 2015;6:1030–1054.

46. Howell GR, Libby RT, Jakobs TC, et al. Axons of retinal ganglion cells are insulted in the optic nerve early in DBA/2J glaucoma. Journal of Cell Biology 2007;179:1523–1537.

47. Johnson EC, Jia L, Cepurna WO, Doser TA, Morrison JC. Global Changes in Optic Nerve Head Gene Expression after Exposure to Elevated Intraocular Pressure in a Rat Glaucoma Model. Investigative Ophthalmology & Visual Science 2007;48:3161–3177.

48. Oikawa K, Teixeira LBC, Keikhosravi A, Eliceiri KW, McLellan GJ. Microstructure and resident cell-types of the feline optic nerve head resemble that of humans. Experimental Eye Research 2021;202:108315.

49. Albon J, Farrant S, Akhtar S, et al. Connective Tissue Structure of the Tree Shrew Optic Nerve and Associated Ageing Changes. Investigative Ophthalmology & Visual Science 2007;48:2134–2144.

50. Chan G, Morgan WH, Yu D-Y, Balaratnasingam C. Quantitative analysis of astrocyte and axonal density relationships: Glia to neuron ratio in the optic nerve laminar regions. Experimental Eye Research 2020;198:108154.

51. Fitzgibbon T, Taylor SF. Retinotopy of the human retinal nerve fibre layer and optic nerve head. J Comp Neurol 1996;375:238–251.

52. Waxman S, Quinn M, Donahue C, et al. Individual astrocyte morphology in the collagenous lamina cribrosa revealed by multicolor DiOlistic labeling. Experimental Eye Research 2023;230:109458.

53. Schlamp CL, Li Y, Dietz JA, Janssen KT, Nickells RW. Progressive ganglion cell loss and optic nerve degeneration in DBA/2J mice is variable and asymmetric. BMC Neurosci 2006;7:66.

54. Kruger K, Tam AS, Lu C, Sretavan DW. Retinal ganglion cell axon progression from the optic chiasm to initiate optic tract development requires cell autonomous function of GAP-43. J Neurosci 1998;18:5692–5705.

55. Ogden TE. Nerve fiber layer of the macaque retina: retinotopic organization. Investigative Ophthalmology & Visual Science 1983;24:85–98.

56. Dichtl A, Jonas JB, Naumann GOH. Course of the optic nerve fibers through the lamina cibrosa in human eyes. Graefe’s Archive for Clinical and Experimental Ophthalmology 1996;234:581–585.

57. Ogden TE. Nerve fiber layer of the owl monkey retina: retinotopic organization. Investigative Ophthalmology & Visual Science 1983;24:265–269.

58. Minckler DS. The Organization of Nerve Fiber Bundles in the Primate Optic Nerve Head. Archives of Ophthalmology 1980;98:1630–1636.

59. Morgan JE, Jeffery G, Foss AJE. Axon deviation in the human lamina cribrosa. British Journal of Ophthalmology 1998;82:680–683.

60. Ritch MD, Hannon BG, Read AT, et al. AxoNet: A deep learning-based tool to count retinal ganglion cell axons. Scientific Reports 2020;10:8034.

61. Reynaud J, Cull G, Wang L, et al. Automated Quantification of Optic Nerve Axons in Primate Glaucomatous and Normal Eyes—Method and Comparison to Semi-Automated Manual Quantification. Investigative Ophthalmology & Visual Science 2012;53:2951–2959.

62. Zarei K, Scheetz TE, Christopher M, et al. Automated Axon Counting in Rodent Optic Nerve Sections with AxonJ. Scientific Reports 2016;6:26559.

63. Hoyt WF, Luis O. Visual Fiber Anatomy in the Infrageniculate Pathway of the Primate: Uncrossed and Crossed Retinal Quadrant Fiber Projections Studied with Nauta Silver Stain. Archives of Ophthalmology 1962;68:94–106.

64. Bray ER, Noga M, Thakor K, et al. 3D Visualization of Individual Regenerating Retinal Ganglion Cell Axons Reveals Surprisingly Complex Growth Paths. eneuro 2017;4:ENEURO.0093-0017.2017.

65. Girkin CA, Fazio MA, Yang H, et al. Variation in the Three-Dimensional Histomorphometry of the Normal Human Optic Nerve Head With Age and Race: Lamina Cribrosa and Peripapillary Scleral Thickness and Position. Investigative Ophthalmology & Visual Science 2017;58:3759–3769.

66. Yang H, Downs JC, Girkin C, et al. 3-D Histomorphometry of the Normal and Early Glaucomatous Monkey Optic Nerve Head: Lamina Cribrosa and Peripapillary Scleral Position and Thickness. Investigative Ophthalmology & Visual Science 2007;48:4597–4607.

67. Downs JC, Yang H, Girkin C, et al. Three-Dimensional Histomorphometry of the Normal and Early Glaucomatous Monkey Optic Nerve Head: Neural Canal and Subarachnoid Space Architecture. Investigative Ophthalmology & Visual Science 2007;48:3195–3208.

68. Lockwood H, Reynaud J, Gardiner S, et al. Lamina Cribrosa Microarchitecture in Normal Monkey Eyes Part 1: Methods and Initial Results. Investigative Ophthalmology & Visual Science 2015;56:1618–1637.

69. Yang H, Downs JC, Sigal IA, Roberts MD, Thompson H, Burgoyne CF. Deformation of the Normal Monkey Optic Nerve Head Connective Tissue after Acute IOP Elevation within 3-D Histomorphometric Reconstructions. Investigative Ophthalmology & Visual Science 2009;50:5785–5799.

70. Roberts MD, Grau V, Grimm J, et al. Remodeling of the Connective Tissue Microarchitecture of the Lamina Cribrosa in Early Experimental Glaucoma. Investigative Ophthalmology & Visual Science 2009;50:681–690.

71. Tran H, Jan N-J, Hu D, et al. Formalin Fixation and Cryosectioning Cause Only Minimal Changes in Shape or Size of Ocular Tissues. Scientific Reports 2017;7:12065.

72. Jones HJ, Girard MJ, White N, et al. Quantitative analysis of three-dimensional fibrillar collagen microstructure within the normal, aged and glaucomatous human optic nerve head. Journal of The Royal Society Interface 2015;12:20150066.

73. Ivers KM, Li C, Patel N, et al. Reproducibility of Measuring Lamina Cribrosa Pore Geometry in Human and Nonhuman Primates with In Vivo Adaptive Optics Imaging. Investigative Ophthalmology & Visual Science 2011;52:5473–5480.

74. Jan N-J, Gomez C, Moed S, et al. Microstructural Crimp of the Lamina Cribrosa and Peripapillary Sclera Collagen Fibers. Investigative Ophthalmology & Visual Science 2017;58:3378–3388.

75. Jan N-J, Lee P-Y, Wallace J, et al. Stretch-Induced Uncrimping of Equatorial Sclera Collagen Bundles. Journal of Biomechanical Engineering 2022;145.

76. Jan N-J, Sigal IA. Collagen fiber recruitment: A microstructural basis for the nonlinear response of the posterior pole of the eye to increases in intraocular pressure. Acta Biomaterialia 2018;72:295–305.

77. Voorhees AP, Jan NJ, Sigal IA. Effects of collagen microstructure and material properties on the deformation of the neural tissues of the lamina cribrosa. Acta Biomaterialia 2017;58:278–290.

78. Zhang L, Albon J, Jones H, et al. Collagen Microstructural Factors Influencing Optic Nerve Head Biomechanics. Investigative Ophthalmology & Visual Science 2015;56:2031–2042.

79. Karimi A, Rahmati SM, Grytz RG, Girkin CA, Downs JC. Modeling the biomechanics of the lamina cribrosa microstructure in the human eye. Acta Biomaterialia 2021;134:357–378.

80. Hua Y, Lu Y, Walker J, et al. Eye-specific 3D modeling of factors influencing oxygen concentration in the lamina cribrosa. Experimental Eye Research 2022;220:109105.

81. Kang MH, Suo M, Balaratnasingam C, Yu PK, Morgan WH, Yu D-Y. Microvascular Density Is Associated With Retinal Ganglion Cell Axonal Volume in the Laminar Compartments of the Human Optic Nerve Head. Investigative Ophthalmology & Visual Science 2018;59:1562–1570.

82. Guan C, Pease ME, Quillen S, et al. Quantitative Microstructural Analysis of Cellular and Tissue Remodeling in Human Glaucoma Optic Nerve Head. Investigative Ophthalmology & Visual Science 2022;63:18–18.

