## supplementary for "Proposing a methodology for axon-centric analysis of IOP-induced mechanical insult"

**S.1. On the use of fluid mechanics to approximate axonal paths**

Our use of fluid mechanics to estimate axonal paths was not intended to imply that the axons are a fluid flowing from inlet to outlet. They are not. The fluid parameters should also not be thought as representing a specific fluid in the axons or similar. Our use of fluid mechanics was done based on the idea that fluid mechanics allowed us to incorporate several reasonable assumptions of axon path characteristics. For example, we assumed that the axon paths from the RNFL to the optic nerve are smooth and continuous from the RNFL to the optic nerve (i.e. no discontinuities), that the axons will pass through the LC pores and not through the LC beams, the CRVs or themselves (i.e. no crossings, splits, bifurcations or similar). In this sense the insight was in realizing that the axon path problem we were working to solve shared characteristics with fluid mechanics. This allowed us to “borrow” fluid mechanics tools to obtain a solution that satisfies simultaneously all the requirements defined above.

It is important to think carefully about the fluid properties and boundary conditions to ensure that the results are physiologically sound. Some fluid mechanics properties may lead to invalid outcomes. For instance, supersonic flows with shockwaves will likely not represent the axon paths due to sharp discontinuities in some parameters. Same if we allowed for sources or sinks.

When we were considering using fluid mechanics, we were encouraged by finding in the literature studies in which the axonal paths in the retina were predicted quite accurately using models from fluid mechanics and electrodynamics. (Tan et al. 2018, Airaksinen et al. 2008, Jansonius et al. 2012, Jansonius et al. 2009) In those studies, researchers were able to tune the fluid or field parameters and validate the models initially using fundus photographs, more recently enhanced by high resolution OCT and SLO. Those studies, however, were limited to predictions of axons in the superficial retina, with predictions ending at the optic disc boundary. Our goal in this study required axonal path predictions into the depths of the ONH and through the LC, but we reckoned that the fundamental processes being similar it was reasonable to use similar modeling approaches.

With respect to specific fluid properties used in this study: we determined adequate properties in a preliminary study, choosing properties that produce a smooth static laminar solution to the flow from which to obtain streamlines representative of axons. Altering the mechanical properties of the fluid or the boundary conditions driving the flow will potentially change the flow. In our preliminary tests we observed that it was possible to vary the properties of the fluid, but as long as the flow was kept laminar, the axonal paths produced were very similar, and so were the mechanical insults derived on them. To illustrate this point for this revision we ran another model in which we changed the viscosity from the baseline of 0.002 Pa s to 5e-4 Pa s. Axon-centric strains and axon paths were obtained for both cases and are shown in Supplementary Figures 1 and 2. As can be seen, the differences are minor both in terms of the axonal paths predicted and on the mechanical insult outcomes.

With respect to the inlet and outlet pressures, we would like to make two comments. First, while several pressures may play a role in optic nerve head and axonal mechanics (IOP, CSFp, BP, etc.), the pressures and their roles in axonal formation are not clear. Second, for the axonal path approximation in this work only the difference between the inlet and outlet pressure is important, as this is what drives the flow. As with the fluid mechanics properties, to illustrate that this parameter did not have a substantial effect on the outcomes, we repeated the modeling changing the inlet/outlet pressures from 0 and 10 kPa to 10 kPa and 20 kPa. The results are shown in the Figure below, included in the manuscript as Supplementary Figure 1.

It is likely that other fluid properties would produce different axonal path predictions. How exactly the paths depend on the fluid properties and the extent of the variations remains unclear and should be the object of a careful and systematic study beyond the scope of this first description of the method. Interestingly, we argue that the fact that fluid properties could change axonal paths predictions presents a valuable opportunity: it should be possible to vary the fluid properties to obtain different axon path predictions that can then be compared with experiments to optimize the accuracy of the path predictions. Unfortunately, to the best of our knowledge at this moment there are still no techniques that can provide detailed experimental maps of the axonal paths through the ONH. Some techniques can reveal paths of a few axons, but to the best of our knowledge none can produce the comprehensive maps needed for optimization. As the experimental techniques are refined, and the data becomes available, it will be possible to re-evaluate axonal paths. We posit that the axonal paths predicted by our method, while imperfect, are a valuable first approximation. The paths obtained in this work seem quite reasonable and consistent with the general idea of axonal paths we have seen in the literature.

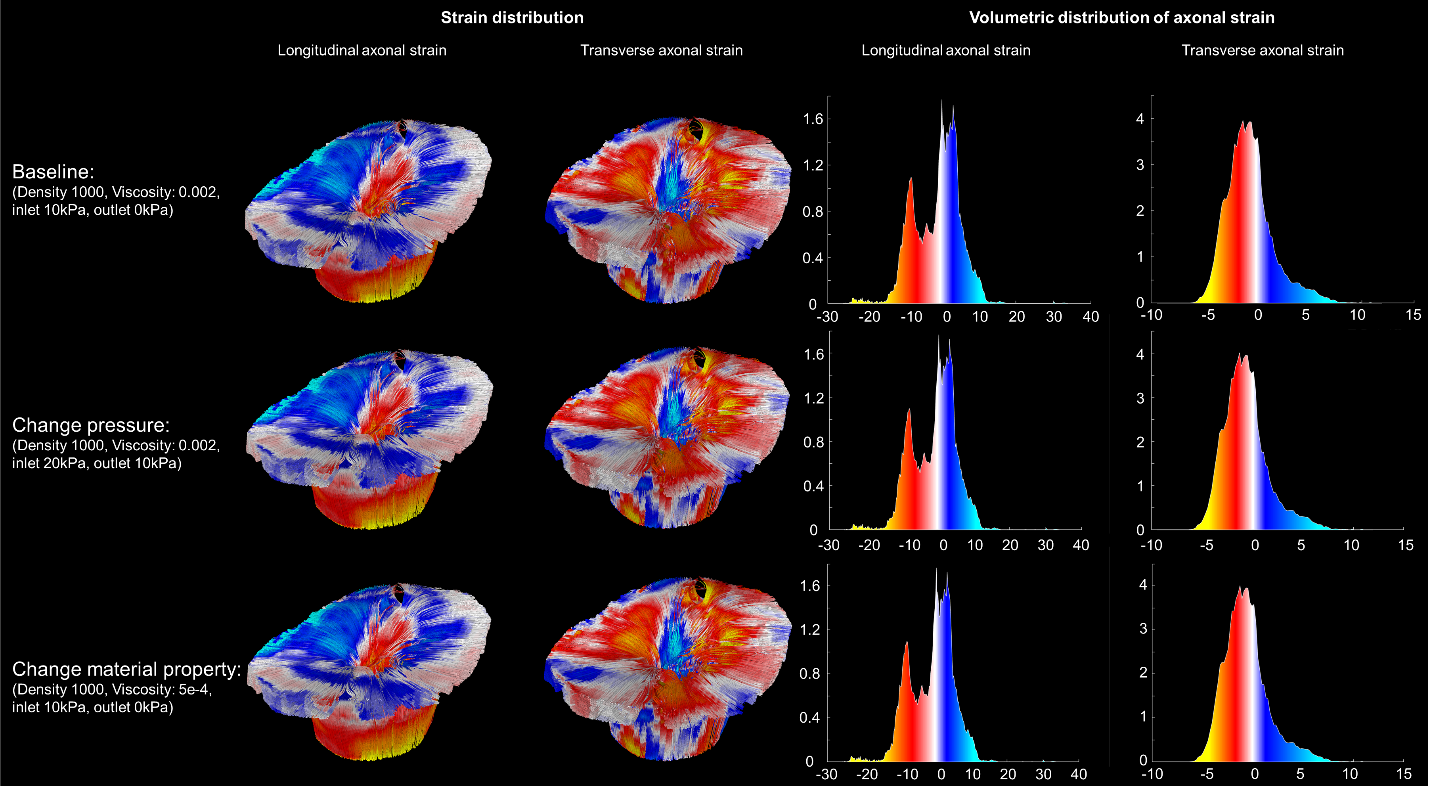

**Supplementary Figure 1.** To evaluate the effects of changes in fluid mechanics parameters on the mechanical insult we compared three cases: 1) baseline: Density 1000, viscosity 0.002, inlet 10 kPa, outlet 0 kPa; 2) change pressures to inlet 20 kPa and outlet 10 kPa; 3) Change fluid viscosity to 5e-4. For each of the cases we show on the left side views of the axons colored by longitudinal and transverse strains, and on the right the distributions of those strains over the axonal volume. All three cases produced essentially the same outcomes.

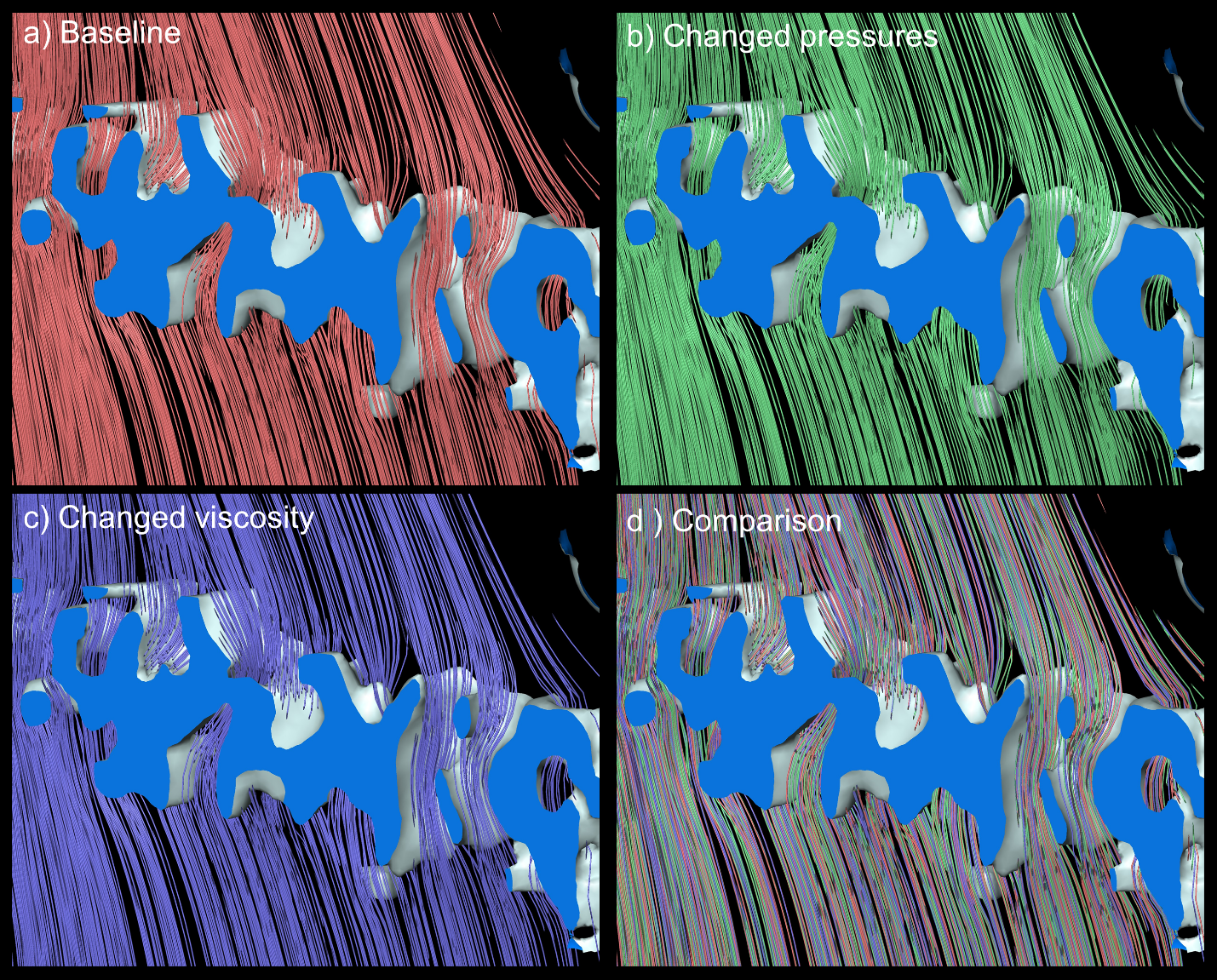

**Supplementary Figure 2.** Effects of changes in fluid mechanics parameters on the predicted axonal paths. Three cases were considered, as shown in the Supplementary Figure 1: **a)** baseline, **b)** change in pressure boundary conditions, and **c)** change in fluid viscosity. The figures show a close-up region of the LC with lines indicating axon paths predicted for the three cases, and **d)** all three overlaid for comparison. In all cases the axons can be seen passing through the LC pores following smooth continuous paths. Interestingly, the paths also result in interspersed regions of higher and lower axonal densities, reminiscent of axonal bundling separated by glia and/or septa. Note that the axons are continuous from RNFL to ON. Any line ends occur because the axons reach the edge of the region selected for view delimited by perpendicular flat faces.

**S.2 Mesh used for the fluid mechanics**

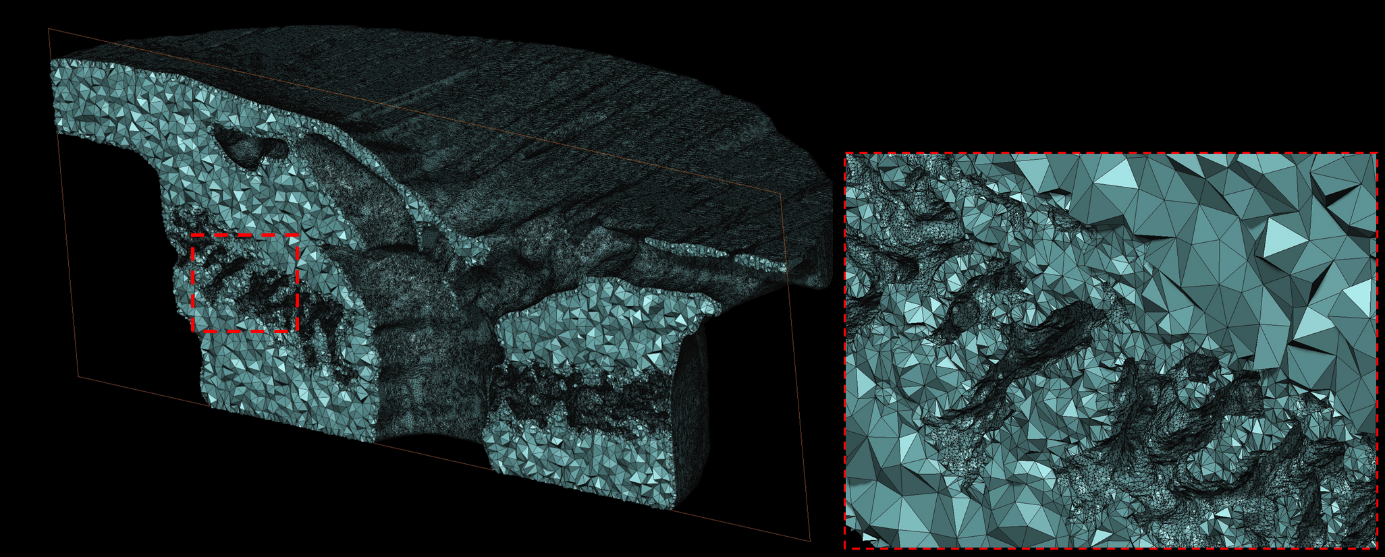

**Supplementary Figure 3.** The mesh of the RGC volume from a cut view.

**S.3 Software and governing equations of CFD simulation**

We used the ABAQUS 6.14 for the CFD calculation. The governing equation is:

$$\rho\frac{\partial\boldsymbol{u}}{\partial t}-\eta\nabla^{2}\boldsymbol{u}+\rho\left( \boldsymbol{u}\cdot\nabla\right)\boldsymbol{u}=F-\nabla p$$

$$\nabla\cdot\boldsymbol{u=}0$$

where $\rho$ is the fluid density (kg/m^3^), $\eta$ (Pa·s) is the fluid viscosity, $\boldsymbol{u}$ is the fluid velocity field (m/s), $p$ is the fluid pressure (Pa), and $F$ is the volume force (N/m^3^). We assumed a zero volume force, and a steady-state condition. The boundary conditions were:

$p=10000$ on ${\partial\Omega}_{in}$

$p=0$ on ${\partial\Omega}_{out}$

$\boldsymbol{u}^{\boldsymbol{*}}\boldsymbol{=}0$ on ${\partial\Omega}_{wall}$

Where ${\partial\Omega}_{in}$ is the inlet surface, ${\partial\Omega}_{out}$ is the outlet surface, ${\partial\Omega}_{wall}$ is the other surfaces. We will provide the full parameters in the supplementary material.

**S.4 Estimated axon paths overlaid on the same OCT**

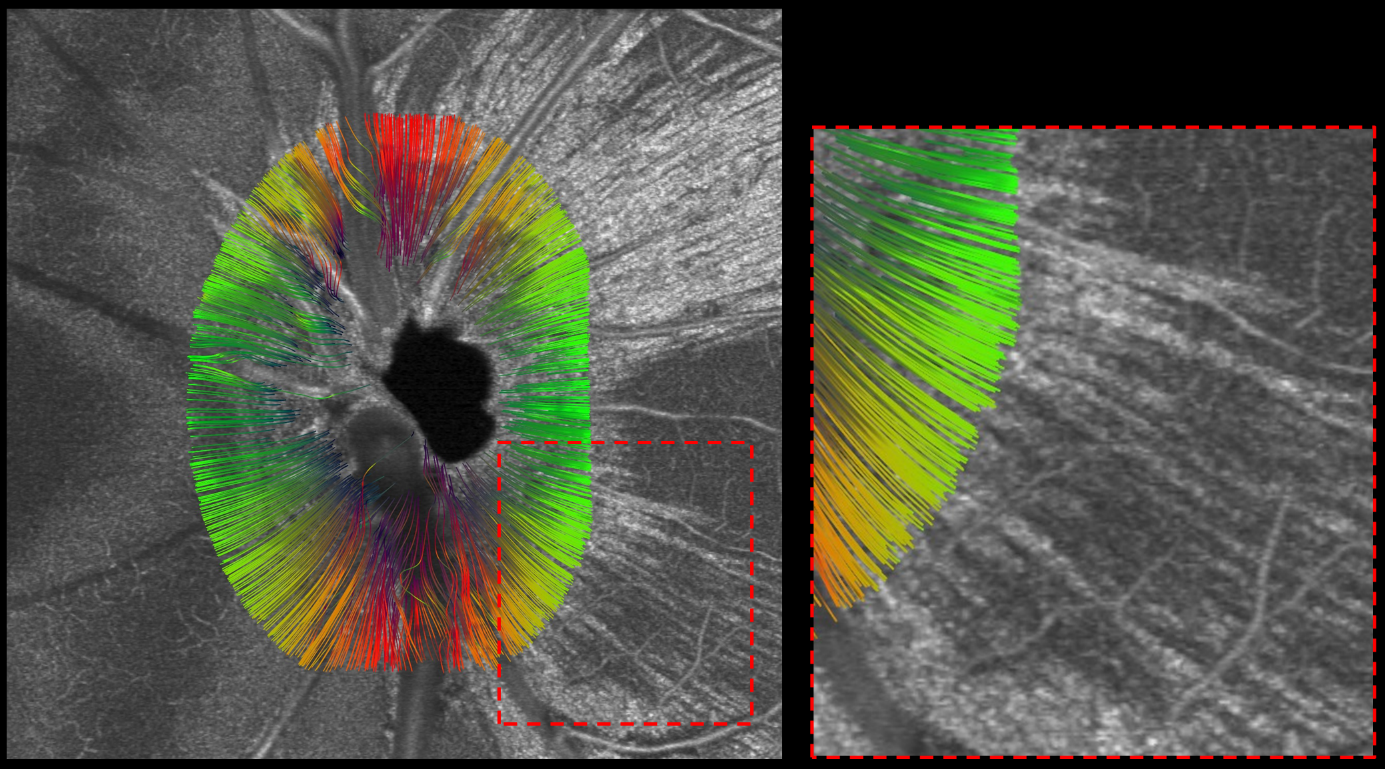

**Supplementary Figure 4.** Coronal en-face view of the NHP ONH. Shown in greyscale is a c-mode virtual cross section through the OCT volume used for estimating the axonal paths. Axonal bundles can be discerned, in this section particularly from about 12 o’clock to 3 o’clock and from 4 o’clock to 5 o’clock. Other clock hours are discernible in more anterior/posterior cross-sections, not shown. Overlaid on the OCT image is shown the axonal paths predicted by our technique, colored by orientation, green horizontal (nasal-temporal) and red vertical (superior-inferior). It is possible to discern that the predicted axonal paths are in reasonable good agreement with the discernible paths. This is more clearly visible in the close-up image (bottom-right). This type of information can be used in the future to optimize the axonal path predictions for a specific eye. Note also that while the OCT image is flat, the axonal paths are 3D, and therefore some are seen “passing over” or “around” blood vessels, at 9 and 11 o’clock.

**S.5 Axonal strain calculation**

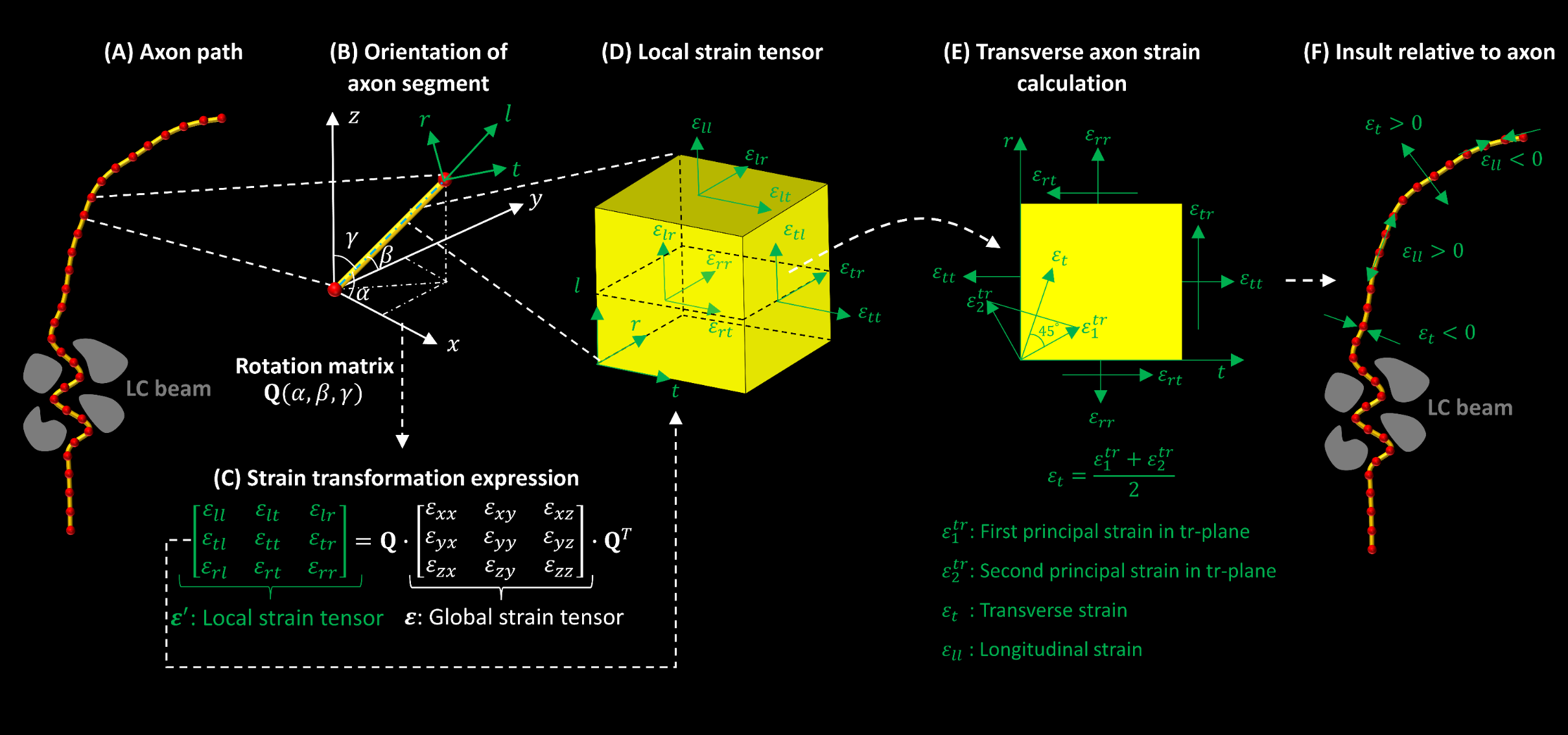

**Supplementary Figure 5** Schematic of steps involved in the calculation of insult relative to the axon. (A) Example of an axon path discretized with points represented by red circles. (B) Zoomed-in view of an axon segment to illustrate local orientation using direction cosines $\alpha, \beta$and $\gamma$ with respect to global cartesian coordinate system $x, y$and $z$ (in white color). The local cartesian coordinate system $l, t$and $r$ (green) where $l$represents direction longitudinal to axon segment, and $t$and $r$represent direction transverse to axon segment. For each segment, the information of directional cosines was used for calculating a rotation matrix. (C) Expression for transforming strain tensor from global ($x, y, z$) to a local coordinate system ($l, t, r$) using a rotation matrix. (D) Schematic of local strain tensor. The strain longitudinal to the axon, represented by $\varepsilon_{ll}$, was extracted directly from the local strain tensor. (E) Schematic of the plane transverse to the axon with local strain $\varepsilon_{tt}, \varepsilon_{rr}$and $\varepsilon_{tr}$ used for calculating transverse insult. We calculated first $(\varepsilon_{1}^{tr})$ and second principal strain ($\varepsilon_{2}^{tr}$) using local strain $\varepsilon_{tt}, \varepsilon_{rr}$and $\varepsilon_{tr}$ values. By following the procedure used for calculating 2D octahedral normal strain, we combined calculated values of both principal strains to get a single value of insult transverse to the axon. The state of a point on the axon path can be identified as stretch or compression based on the sign (positive means stretch and negative means compression) of transverse insult. (F) Schematic of relative insult to the axon (longitudinal and transverse) in stretch (value greater than zero) and compression (value less than zero).
